# Positive selection drives *cis-*regulatory evolution across the threespine stickleback Y chromosome

**DOI:** 10.1101/2022.11.03.515077

**Authors:** Daniel E. Shaw, Alice Shanfelter Naftaly, Michael A. White

**Author notes:** **Corresponding Author: Michael A. White Department of Genetics, University of Georgia 120 Green St, Athens, GA 30602**.

## Abstract

Allele-specific gene expression evolves rapidly on heteromorphic sex chromosomes. Over time, the accumulation of mutations on the Y chromosome leads to widespread loss of gametolog expression, relative to the X chromosome. It remains unclear if expression evolution on degrading Y chromosomes is primarily driven by mutations that accumulate through processes of selective interference, or if positive selection can also favor the downregulation of coding regions on the Y chromosome that contain deleterious mutations. Identifying the relative rates of *cis*-regulatory sequence evolution across Y chromosomes has been challenging due to the limited number of reference assemblies. The threespine stickleback (*Gasterosteus aculeatus*) Y chromosome is an excellent model to identify how regulatory mutations accumulate on Y chromosomes due to its intermediate state of divergence from the X chromosome. A large number of Y-linked gametologs still exist across three differently aged evolutionary strata to test these hypotheses. We found that putative enhancer regions on the Y chromosome exhibited elevated substitution rates and decreased polymorphism when compared to non-functional sites, like intergenic regions and synonymous sites. This suggests that many *cis*-regulatory regions are under positive selection on the Y chromosome. This divergence was correlated with X-biased gametolog expression, indicating the loss of expression from the Y chromosome may be favored by selection. Our findings provide evidence that Y-linked *cis*-regulatory regions exhibit signs of positive selection quickly after the suppression of recombination and allow comparisons with recent theoretical models that suggest the rapid divergence of regulatory regions may be favored to mask deleterious mutations on the Y chromosome.

## Introduction

The evolution of heteromorphic sex chromosomes has occurred many times among species (Bachtrog 2013; Bachtrog, et al. 2014). Heteromorphic sex chromosomes evolve once recombination is suppressed between the X and Y (or Z and W) (Muller 1918; Charlesworth 1978; Charlesworth and Charlesworth 2000). After recombination is suppressed, the Y chromosome rapidly accumulates mutations, leading to sequence degeneration (Charlesworth and Charlesworth 2000). Empirical evidence of this process has focused on the sequence evolution of coding regions on sex chromosomes. Broad comparative work has revealed that coding sequence evolution can follow one of several different evolutionary trajectories. Although many ancestral Y-linked genes are lost because of sequence degeneration through the accumulation of deleterious mutations, growing evidence indicates not all genes are lost. Some genes appear to be dosage sensitive and are under strong purifying selection to be retained on the Y chromosome (Bellott, et al. 2014; White, et al. 2015; Bellott, et al. 2017; Peichel, et al. 2020). Novel genes with sex-specific functions can also accumulate on sex-limited chromosomes via translocation and subsequent gene duplication (Soh, et al. 2014; Mahajan and Bachtrog 2017; Ellison and Bachtrog 2019; Hughes, et al. 2020; Peichel, et al. 2020; Chang, et al. 2022).

The accumulation of mutations is not restricted to coding sequences and can also occur in *cis*-regulatory regions, leading to changes in expression from the sex-limited chromosome. Gametolog expression (ancestral genes still shared between the X and Y chromosomes) has been shown to evolve rapidly on degenerating sex chromosomes, often resulting in the loss of expression from the sex-limited chromosome (Y and W) (Meisel, et al. 2012a; Muyle, et al. 2012; Ayers, et al. 2013; Singh, et al. 2014; White, et al. 2015; Beaudry, et al. 2017; Muyle, et al. 2018; Rodríguez Lorenzo, et al. 2018; Martin, et al. 2019; Veltsos, et al. 2019; Wei and Bachtrog 2019; Shaw and White 2022). Although deleterious regulatory mutations may accumulate through selective interference (Charlesworth and Charlesworth 2000; Bachtrog 2008), selection may also favor mutations within *cis*-regulatory regions to downregulate coding regions with deleterious mutations (Orr and Kim 1998; Bachtrog 2006). Recent theory supports the role of positive selection driving the rapid accumulation of mutations to downregulate the Y-linked allele and upregulate the X-linked allele to maintain ancestral dosage balance (Lenormand, et al. 2020; Lenormand and Roze 2022).

One challenge to characterizing the molecular evolution of regulatory regions on sex chromosomes is that complete Y chromosome assemblies are only available for a limited number of taxa. Reference assemblies are needed to identify the substitutions that are accumulating within non-coding regions. Additionally, many of the available Y chromosome assemblies are from model organisms that have sex chromosomes that are highly degenerated (Hughes, et al. 2010; Bellott, et al. 2014; Soh, et al. 2014; Tomaszkiewicz, et al. 2016; Mahajan, et al. 2018; Hughes, et al. 2020). These species only have a few remaining ancestral gametologs on the Y chromosome, thus limiting the number of genes available to study how *cis*-regulatory evolution potentially leads to expression differences between the X and Y. Species with recently derived sex chromosomes that still harbor many ancestral Y-linked gametologs at varied stages of degeneration are needed to understand how substitutions within *cis*-regulatory regions lead to the evolution of expression differences between the X and Y.

In addition to having a chromosome-scale assembly of the Y chromosome, an additional challenge is annotating regulatory regions (Reviewed in Shaw & White, 2022). Recent approaches have focused on identifying regions of the genome with accessible chromatin that may contain transcription factor binding sites to regulate gene expression (Ricci, et al. 2019). Chromatin accessibility profiling techniques, like ATAC-seq (Buenrostro, et al. 2013), utilize short-read sequencing to profile accessible chromatin regions (ACRs) at a fine-scale across the genome. Across autosomes, ACRs show signatures of purifying selection relative to other intergenic regions, consistent with these regions containing important functional elements for gene regulation (Connelly, et al. 2014; Lu, et al. 2019; Horvath, et al. 2021). Profiling ACRs on sex chromosomes would therefore provide a means to study the molecular evolution of *cis*-regulatory regions in the context of Y degeneration.

The threespine stickleback fish (*Gasterosteus aculeatus*) is an excellent model to study the evolution of *cis*-regulatory regions on sex chromosomes. Threespine stickleback fish have a high-quality reference assembly of the Y chromosome and this chromosome is more recently derived compared to many other species with available chromosome-scale assemblies (Hughes, et al. 2010; Hughes, et al. 2012; Bellott, et al. 2014; Bellott, et al. 2017; Hughes, et al. 2020; Peichel, et al. 2020). The threespine stickleback sex chromosomes contain multiple evolutionary strata at different stages of degeneration (Roesti, et al. 2013; White, et al. 2015; Peichel, et al. 2020), allowing for comparisons of sequence evolution at different temporal scales. Strata form if recombination is suppressed between sex chromosomes in multiple steps (Bachtrog 2013). Each subsequent phase of recombination suppression is followed by the accumulation of substitutions on the Y chromosome, leading to increased sequence divergence between the X and Y. The age of individual strata can therefore be distinguished by varied levels of synonymous divergence. In threespine stickleback fish, the oldest stratum is estimated to have formed around 22 million years ago. Two younger strata formed between 4 and 6 million years ago (Peichel, et al. 2020; Sardell, et al. 2021). Stratum one has undergone the most extensive degeneration, with only 18% of ancestral coding regions remaining on the Y chromosome (Peichel, et al. 2020). Many of the coding regions of these gametologs that remain in stratum one on the Y chromosome show signatures of purifying selection and are enriched for dosage sensitive functions (White, et al. 2015; Peichel, et al. 2020). Interestingly, chromosome-wide dosage compensation has not evolved to counter the loss of expression for genes where the coding sequence has degenerated on the Y chromosome (White, et al. 2015). The lack of chromosome-wide dosage compensation enables a gene-by-gene comparison of *cis-*regulatory changes to understand how expression of X-linked alleles evolves in response to degenerating coding regions on the Y.

Here, we leveraged ATAC-seq from two different tissues to identify *cis-*regulatory regions on the sex chromosomes and an autosome. We found sex-linked *cis-*regulatory regions exhibited signatures of positive selection, relative to non-functional, intergenic control regions. Our findings complement existing sex chromosome evolution theory that suggests the accumulation of mutations within *cis*-regulatory regions may be beneficial to silence expression from a degenerating Y chromosome.

## Results

### Nucleotide substitutions are accumulating at high rates within accessible chromatin regions

The completion of the threespine stickleback Y chromosome assembly (Peichel, et al. 2020) allowed us to thoroughly examine regulatory evolution among sex-linked gametologs shared between the sex chromosomes. ACRs putatively contain *cis-*regulatory elements that serve as domains for transcription factor binding. High sequence divergence within ACRs therefore represent regions of the genome that are likely experiencing *cis-*regulatory evolution. We first sought to compare divergence of ACRs between the X and Y chromosomes to previously characterized synonymous divergence throughout coding regions. Crossing over was suppressed across the threespine stickleback sex chromosomes in at least three separate events, forming strata with distinct levels of divergence (stratum one: oldest; stratum two: intermediate; stratum three: youngest) (Peichel, et al. 2020). To survey ACRs, we utilized an Assay for Transposase Accessible Chromatin sequencing (ATAC-seq) from liver tissue (Naftaly, et al. 2021) and testis tissue. We defined a set of non-coding ACRs that were likely present in the ancestor of the X and Y chromosomes, by identifying X-linked ACRs within intergenic sequence that still had homologous sequence on the Y chromosome and homologous sequence in the ancestral autosome (homologous sequence in ninespine stickleback chromosome 19). To calculate divergence, we compared these regions with the homologous sequence on the Y chromosome, which may no longer be associated with accessible chromatin. In the liver, we found a total of 1,279 X-linked ACRs. We were able to align 948 (74.1%) of these ACRs to homologous regions on the Y chromosome and the ancestral ninespine stickleback *(Pungititus pungitius)* autosome 19. 869 liver ACRs were within 50 kb of a coding region shared by all three chromosomes. In the testis, we identified 896 X-linked ACRs. Compared to the liver, we found far fewer orthologous regions on the Y chromosome and autosome 19 in the ninespine stickleback (118 total aligned; 13.2%), suggesting *cis*-regulatory regions within testis ACRs may be more rapidly evolving. All 118 of the aligned testis ACRs were within 50 kb of a coding region shared by all three chromosomes. We used these aligned sets of ACRs that were close to conserved coding regions for all subsequent analyses.

We estimated divergence between the X and Y chromosomes among ACRs in each tissue and compared this to synonymous and non-synonymous substitutions within coding regions across the sex chromosomes. Coding regions across all three strata on the threespine stickleback Y chromosome are accumulating non-synonymous substitutions at a higher rate than autosomes due to a reduced efficacy of selection (White, et al. 2015; Peichel, et al. 2020). We first compared the divergence among testis and liver ACRs to non-synonymous substitution rates within coding regions to determine if mutations were accumulating at a similar rate. We found liver and testis ACRs have much higher divergence compared to non-synonymous mutations among all three strata (Figure 1) (P < 0.001; Kruskal-Wallis and Dunn’s Test). We also compared ACR divergence with synonymous substitutions, which is used as measure of neutral evolution. Testis ACRs had divergence higher than neutral, synonymous substitutions in all three strata (Figure 1) (P < 0.001; Kruskal-Wallis and Dunn’s Test). Liver ACR’s exhibited divergence that was higher than synonymous substitutions in the two oldest strata (one and two) (P < 0.001; Kruskal-Wallis and Dunn’s Test) (Figure 1), indicating these regions are also evolving rapidly. In the youngest stratum (three), the liver ACRs were not significantly different than synonymous substitutions (P = 0.413, Kruskal-Wallis and Dunn’s Test). Additionally, ACR divergence was distinct between all three evolutionary strata (P < 0.001; Kruskal-Wallis and Dunn’s test) and corresponded with the extent of evolutionary divergence of each stratum. ACR divergence higher than both non-synonymous and synonymous substitutions strongly suggests *cis-*regulatory regions are evolving rapidly across all three strata. However, this analysis alone cannot rule out the possibility that purifying selection may still be operating within coding regions on the Y chromosome, reducing non-synonymous substitutions. Synonymous substitutions may also be under weak purifying selection (i.e. codon bias) (Bachtrog 2005; Lawrie, et al. 2013). In this scenario, we may observe higher divergence rates within ACRs relative to coding regions.

**Figure 1.**
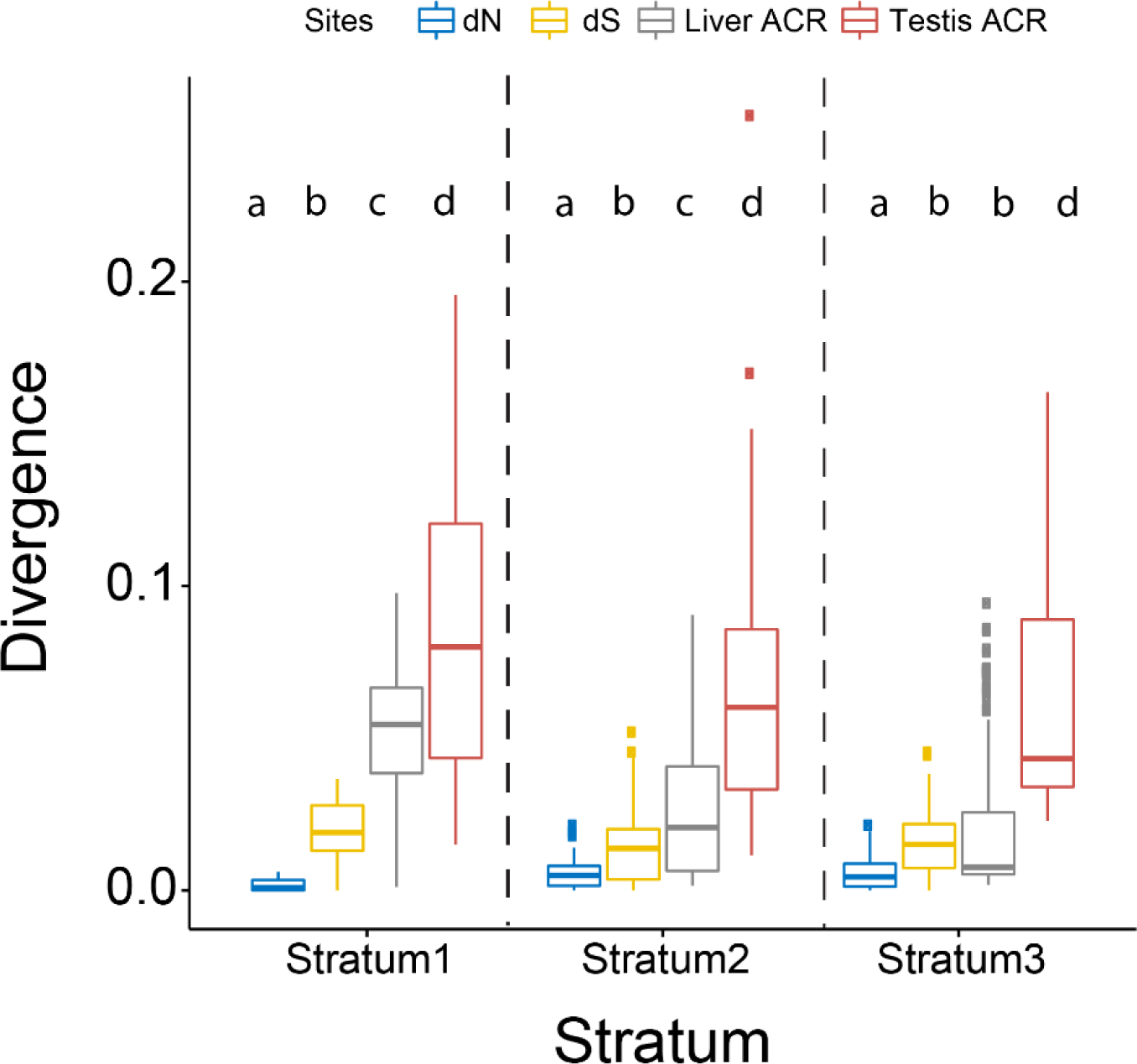
XY divergence within accessible chromatin regions and coding regions. Accessible chromatin regions (ACRs) show elevated divergence between the X and Y, relative to nonsynonymous and synonymous substitutions in coding regions. Protein-coding divergence was estimated between the X and Y chromosomes at all shared gametologs. ACRs were identified on the X chromosome and mapped to the Y chromosome. Substitutions within ACRs were summed and divided by the total length of the ACR to calculate divergence. Stratum one: 12 gametologs, 442 liver ACRs, 30 testis ACRs; Stratum two: 71 gametologs, 228 liver ACRs, 37 testis ACRs; Stratum three: 46 gametologs, 199 liver ACRs 51 testis ACRs. Rates of sequence divergence for all 4 categories were compared using a Kruskal-Wallis test for each stratum (P < 0.001 for all strata). A post-hoc Dunn’s test was used to identify which groups were significantly different from one another. P-values were adjusted for multiple comparisons with a Bonferroni correction. The letters above each plot indicate the significant differences between sites within each stratum (P < 0.001). Box plots that share a letter assignment are not significantly different from one another (P > 0.001).

We therefore sought to compare ACR divergence with non-functional regions throughout the sex chromosomes as well as ACR divergence throughout a representative autosome. We used the orthologous sequence from the ninespine stickleback fish to infer substitution rates throughout the X and Y chromosomes separately as well as throughout autosome 18. We defined non-functional control regions as intergenic sequence of the chromosomes that did not overlap known gene annotations, repetitive elements, or our previously defined ACRs (see methods). Although our ability to define non-functional regions is limited to the currently available annotations, this represents our best approximation of regions of the genome that are not under selection. If ACRs are evolving faster than expected under neutrality, we would predict the average ACR sequence divergence would exceed the rate of divergence observed across permutations of non-functional control regions (Figure 2A). However, if ACR divergence is similar to the control regions (Figure 2A), this would suggest substitutions are accumulating in ACRs at the same rate as the remainder of the Y chromosome, which is subject to sequence degeneration across the non-crossover region (Charlesworth and Charlesworth 2000).

**Figure 2.**
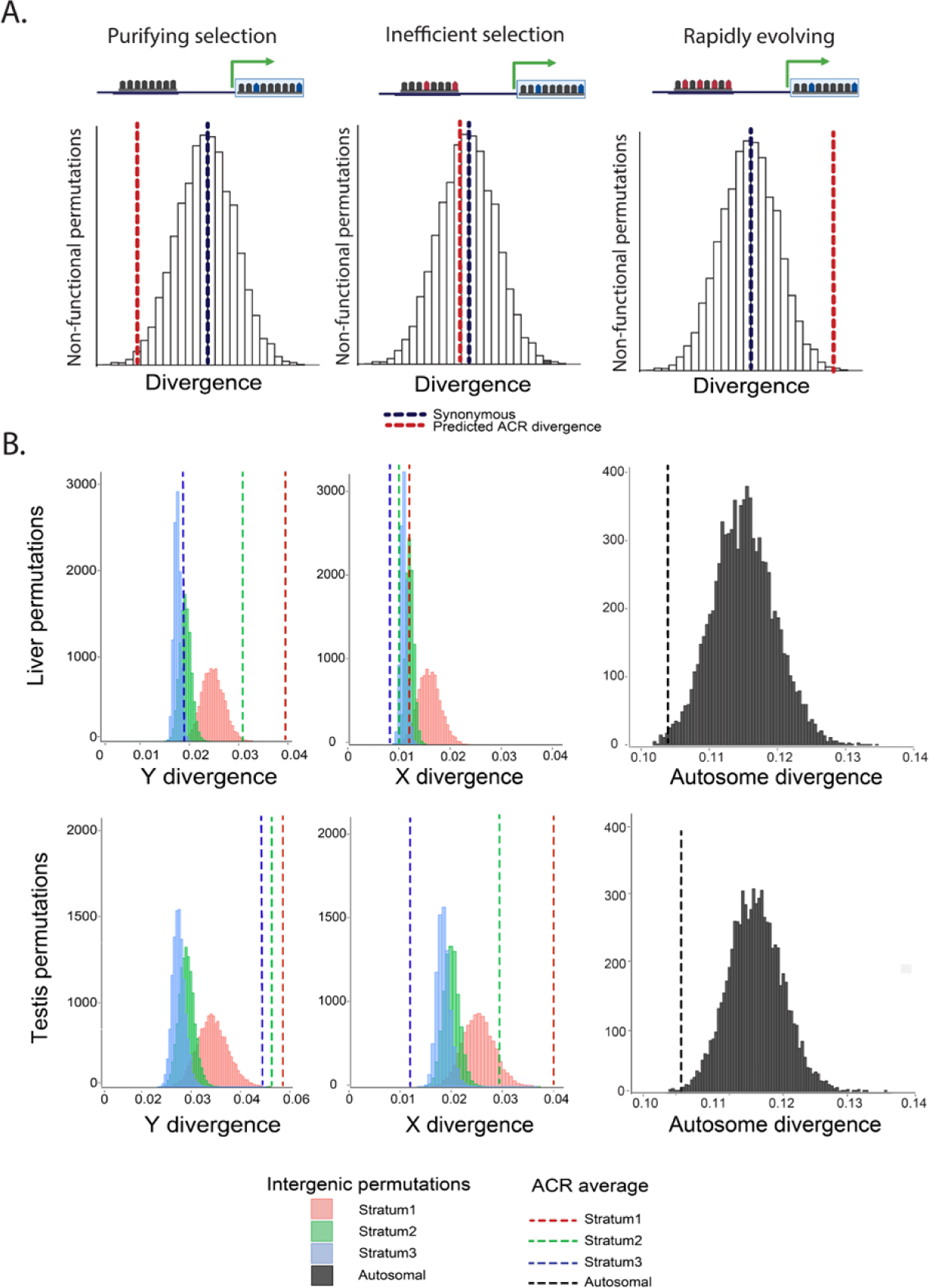
Accessible chromatin divergence between threespine and ninespine stickleback. **A.** Predicted substitution rates within accessible regions. ACR substitution rates (red) lower than random non-functional control regions (white) and synonymous substitutions (blue) would indicate purifying selection (left). ACR substitution rates roughly equal to non-functional control regions would indicate mutations are accumulating due to inefficient selection from the loss of recombination (middle). ACR substitution rates higher than non-functional control regions would indicate positive selection. **B.** Y-linked ACR divergence is elevated over non-functional intergenic regions. X and Y ACR divergence was compared to randomly drawn non-functional regions matched for GC content (10,000 permutations). The average ACR divergence of each stratum is shown by the dotted line (stratum one: pink, stratum two: green, and stratum three: blue). ACRs were identified on the X chromosome and mapped to the Y chromosome and homologous ninespine stickleback autosome 19 from liver (top) and testis (bottom). Autosome divergence is from chromosome 18. For the sex chromosomes, we identified variants that were unique to the X and Y, by aligning to the outgroup and ignoring fixed differences between species that likely occurred before the sex chromosomes evolved. X- and Y-specific variants within ACRs were summed and divided by the total length of the ACR to calculate divergence. Stratum one: 442 liver ACRs and 30 testis ACRs; stratum two: 228 liver ACRs and 37 testis ACRs; stratum three: 129 liver ACRs and 46 testis ACRs; autosomal: 1056 liver ACRs and 305 testis ACRs. Graphic made with BioRender.com

ACRs functional in the liver exhibited higher substitution rates on the Y chromosome than on the X chromosome. Y-linked ACRs had significantly higher substitution rates compared to the non-functional control regions for the two oldest evolutionary strata (Figure 2B; P < 0.001; stratum one and two; 10,000 permutations). In contrast to the Y-linked ACRs, we found the X-linked liver ACR sequence divergence was significantly lower than non-functional control regions (Figure 2B; Stratum one: P = 0.016; Stratum two and three: P < 0.001; 10,000 permutations). This suggests that ACRs may be functionally constrained more on the X chromosome, compared to Y-linked ACRs. We observed a similar pattern on the Y chromosome for ACRs functional in testis tissue. Y-linked ACRs had significantly higher substitution rates in all three evolutionary strata, compared to non-functional control regions (Figure 2B; P < 0.001, 10,000 permutations). However, we observed X-linked ACRs from testis tissue, also exhibited elevated nucleotide substitutions in the oldest two evolutionary strata (Figure 2B; P < 0.001; 10,000 permutations), while the youngest stratum was significantly lower than intergenic regions (Figure 2B; P < 0.001; 10,000 permutations). This indicates some *cis*-regulatory elements exhibited accelerated substitution rates on the X chromosome and this occurs in a tissue-specific fashion.

Autosomal ACRs have been shown to be under purifying selection in other species (Horvath, et al. 2021). We therefore tested whether threespine stickleback autosomal ACRs also exhibited purifying selection, similar to the pattern we observed on the X chromosome in liver. Consistent with patterns of purifying selection, we found autosomal ACRs exhibited lower sequence divergence compared to non-functional control regions in both liver (Figure 2B; P = 0.002; 10,000 permutations) and testis tissue (Figure 2B; P < 0.001; 10,000 permutations).

### Accessible chromatin regions exhibit signatures of selection

An excess of substitutions on the Y chromosome could be caused by relaxed purifying selection or positive selection. To distinguish between these alternatives, we used short-read sequencing of 12 males from a freshwater population of stickleback fish (Shanfelter, et al. 2019) to search for signatures of selection within ACRs. In the absence of selection, the ratio of polymorphism to sequence divergence should be equal between functional and non-functional sites. Deviation from this expectation would imply these regions are under selection. To explore this, we performed a modified McDonald-Kreitman (MK) test on non-coding sequences (Andolfatto 2005). We compared the number of X- and Y-specific substitutions to the number of polymorphisms segregating within a population for ACRs and non-functional control sequences. Except for stratum two on the X chromosome, sex chromosome ACRs from the liver and the testis had a greater proportion of sites fixed by positive selection (*α*), compared to autosomal ACRs (P < 0.001; chi-square test) (Table 1). We also identified the direction of selection acting on ACRs by calculating the odds-ratio of the McDonald Kreitman test, known as the neutrality index (NI) (Rand and Kann 1996). Liver ACRs of the sex chromosomes had an NI < 1 (P < 0.001; chi-square test), consistent with positive selection (Figure 3; Table 1). The only exception we found was for one of the younger strata on the X chromosome (stratum two), which had an elevated NI consistent with purifying selection. Despite having a mean divergence lower than intergenic sequence, we found signatures of positive selection on the X chromosome for strata one and three. In these cases, the departure from neutrality was driven by the low number of polymorphisms within ACRs compared to intergenic sequence, rather than high divergence within ACRs (Table 1). In these cases, the reduced polymorphism may be due to recent positive selection (Stoletzki and Eyre-Walker 2010, Hughes, et al. 2008). Each stratum on the Y chromosome had a clear signature of positive selection (NI < 1; P < 0.001). Compared to the X chromosome and the autosome, the Y chromosome liver ACRs had stronger signatures of positive selection, indicative by a non-overlapping bootstrap interval. For the testis ACRs, NI was indistinguishable between the X and Y chromosome, and both sex chromosomes had non-overlapping bootstrap intervals with the representative autosome, indicating stronger positive selection acting on sex-linked ACRs. Similar to X-linked ACRs in liver, we found that testis ACRs in stratum three on the X chromosome exhibited an NI < 1, due to a low number of polymorphisms in ACR rather than an excess in divergence.

**Figure 3.**
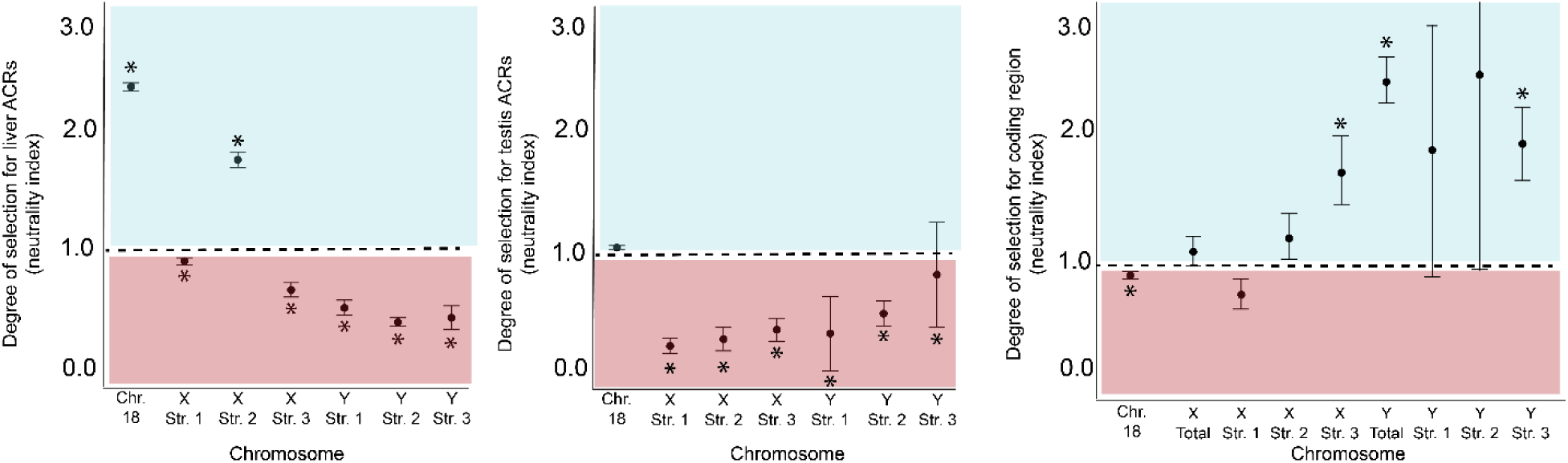
Quantifying adaptive divergence and departure from neutrality among accessible chromatin regions and coding regions. Sex-linked ACRs exhibited signatures of positive selection in both liver (left) and testis (middle). Sex-linked coding regions (right) are mostly under purifying selection or not significantly different than neutral expectations. The neutrality index (NI) was calculated using the odds ratio of the McDonald Kreitman test. NI > 1 reflects purifying selection (blue) while NI < 1 is a signature of positive selection (red). Asterisks denote significant departures from neutrality determined by a chi-square test. Error bars represent two standard deviations calculated from 10,000 bootstrap replicates.

**Table 1.** McDonald-Kreitman Test for non-coding ACRs.

| Chromosome | Stratum | Tissue | D(ACR) | D(INT) | P(ACR) | P(INT) | $\alpha$ | NI | ChiSquare | P-value |
| --- | --- | --- | --- | --- | --- | --- | --- | --- | --- | --- |
| Y | Stratum 1 | Testis | 739 | 555 | 18 | 46 | 0.706 | 0.294 | 20.77 | P<0.001 |
| Y | Stratum 2 | Testis | 1292 | 912 | 2 | 10 | 0.859 | 0.141 | 8.65 | P=0.003 |
| Y | Stratum 3 | Testis | 454 | 194 | 7 | 16 | 0.813 | 0.187 | 14.43 | P<0.001 |
| Y | Stratum 1 | Liver | 89313 | 52716 | 243 | 923 | 0.845 | 0.155 | 870.81 | P<0.001 |
| Y | Stratum 2 | Liver | 30619 | 20743 | 210 | 388 | 0.633 | 0.367 | 147.01 | P<0.001 |
| Y | Stratum 3 | Liver | 16889 | 16875 | 321 | 412 | 0.222 | 0.778 | 11.13 | P<0.001 |
| X | Stratum 1 | Testis | 878 | 760 | 452 | 18109 | 0.978 | 0.022 | 6406.49 | P<0.001 |
| X | Stratum 2 | Testis | 1225 | 1037 | 52 | 460 | 0.904 | 0.096 | 325.35 | P<0.001 |
| X | Stratum 3 | Testis | 382 | 302 | 91 | 365 | 0.803 | 0.197 | 145.19 | P<0.001 |
| X | Stratum 1 | Liver | 26658 | 34849 | 6009 | 10003 | 0.215 | 0.785 | 176 | P<0.001 |
| X | Stratum 2 | Liver | 7121 | 13850 | 1265 | 1321 | -0.863 | 1.863 | 224.76 | P<0.001 |
| X | Stratum 3 | Liver | 9644 | 13485 | 2618 | 7618 | 0.519 | 0.481 | 793.25 | P<0.001 |
| Chr18 | NA | Liver | 23380 | 25607 | 9533 | 4193 | -1.49 | 2.49 | 2028.33 | P<0.001 |
| Chr18 | NA | Testis | 8715 | 9545 | 1509 | 1563 | -0.057 | 1.057 | 2.05 | P=0.153 |

We compared the patterns we observed across ACRs with coding regions. We found that coding regions across the entire Y chromosome had an excess of non-synonymous polymorphisms (Figure 3; NI > 1; P < 0.001; chi-square test), consistent with relaxed purifying selection on a degenerating Y chromosome. This pattern was largely driven by the youngest two strata of the Y chromosome (Table 2). The oldest stratum (one) did not deviate from neutral expectations (P = 0.112). Coding regions across the X chromosome were not significantly different than neutral expectations (NI = 1; P = 0.281; chi-square test). When split by evolutionary strata only the youngest stratum (three) deviated from neutral expectations after correcting for multiple comparisons. In stratum three, the coding regions exhibited purifying selection (NI > 1; Pn < Ps; P < 0.001; chi-square test). Unlike the Y chromosome, the X entire chromosome did not have an excess of non-synonymous polymorphism (Table 2). A gene-by-gene MK test revealed that 50% of X-linked gametologs had an NI > 1, compared to only 25% that had an excess of amino acid substitutions (Supplemental material). Importantly, sex-linked coding regions were not under widespread positive selection, like ACRs.

**Table 2.**
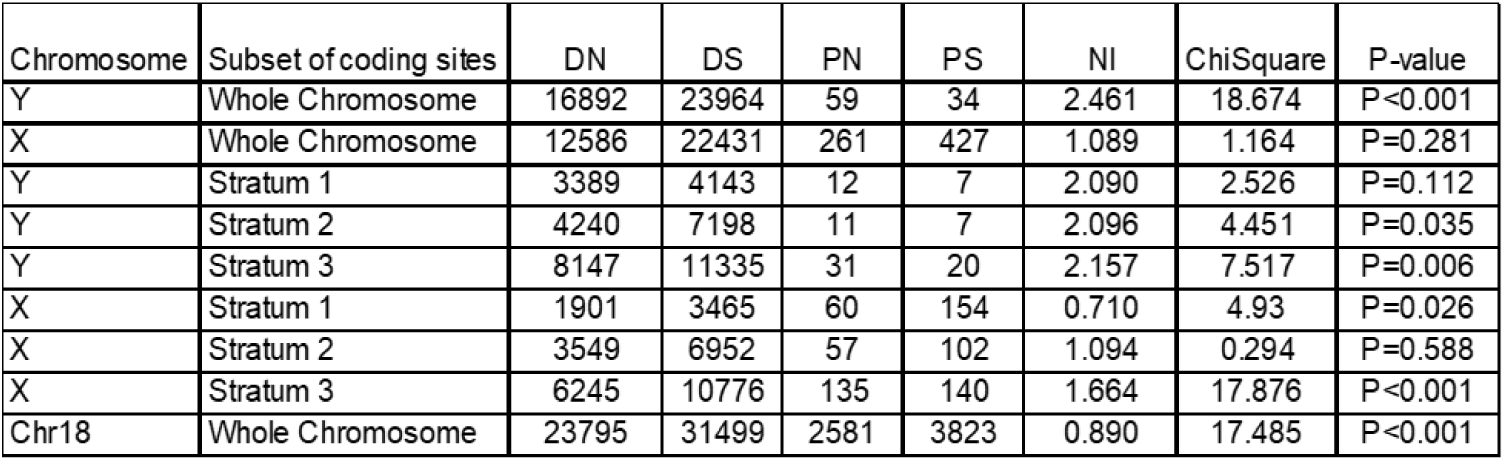
McDonald-Kreitman Test for coding regions.

### Reduced polymorphism throughout the sex chromosomes

Due to a lower effective population size, sex chromosomes are expected to have lower nucleotide diversity relative to the autosomes, given an equal sex ratio in the population. To test whether nucleotide diversity is reduced on the threespine stickleback sex chromosomes, we estimated average nucleotide diversity in sliding windows across the sex chromosomes and autosome 18. Across windows, we found Y chromosome nucleotide diversity was much lower than the expected 25% of nucleotide diversity compared to autosomes (Y chromosome pi: 0.0001; autosome pi: 0.0043; Supplemental Figure 1). This suggests that selection acting on linked deleterious or beneficial mutations has lowered nucleotide diversity beyond neutral expectations on the Y chromosome. The X chromosome is predicted to contain 75% of the nucleotide diversity compared to the autosomes. We found that the non-PAR regions of the X chromosome had 45% of the amount of nucleotide diversity as autosomes, indicating the X chromosome may also be under selection, reducing nucleotide diversity beyond neutral expectations (non-PAR X chromosome pi: 0.0019; autosome pi: 0.0043; Supplemental Figure 1). We found that the Y chromosome had consistent signatures of low nucleotide diversity across all strata (Supplemental Figure 1). However, there were distinct differences in nucleotide diversity for each stratum and the PAR on the X chromosome (Supplemental Figure 1; P < 0.001 for all comparisons; Kruskal-Wallis and Dunn’s Test), suggesting each region of the X chromosome may be under different selection pressures depending on gene and regulatory content.

### Y-linked substitutions are correlated with biased expression from the X chromosome

Selection should favor the loss of expression from the Y chromosome in order to silence coding regions that have accumulated deleterious mutations (Orr and Kim 1998; Bachtrog 2006; Lenormand, et al. 2020). To test if Y-linked *cis*-regulatory divergence was associated with changes in expression, we compared ACR sequence divergence to changes in allelic expression on the X and Y from liver and testis RNA-seq transcriptomes (Peichel, et al. 2020). To maximize the total number of genes for this analysis, we pooled genes across all three evolutionary strata. We found that sequence divergence among liver ACRs that were proximal to genes was a predictor of allele-specific expression. We observed higher expression of the X-linked gametolog, relative to the Y-linked gametolog, when ACRs on the Y chromosome had more substitutions (Figure 4A; N = 51; P = 0.004; R=0.4; Spearman’s rank correlation; 95% Confidence Interval via bootstrapping (0.1249, 0.5769)), indicating that ACR divergence explains around 16% of the X-biased expression observed in liver tissue We also observed a correlation between expression and substitutions within ACRs on the X chromosome (Figure 4C; N = 51; P = 0.033; R=0.30; Spearman’s rank correlation; 95% Confidence Interval from bootstrapping (0.0909, 0.4742)). The slopes of each linear regression were indistinguishable between the X and Y chromosome (P =0.590; Fisher Z transformation). Although we do not have gene expression from the ninespine stickleback to determine which sex chromosome is deviating from the ancestral expression level, the Y-linked substitutions indicate the X-biased expression we observe is likely due to downregulating the Y chromosome.

**Figure 4.**
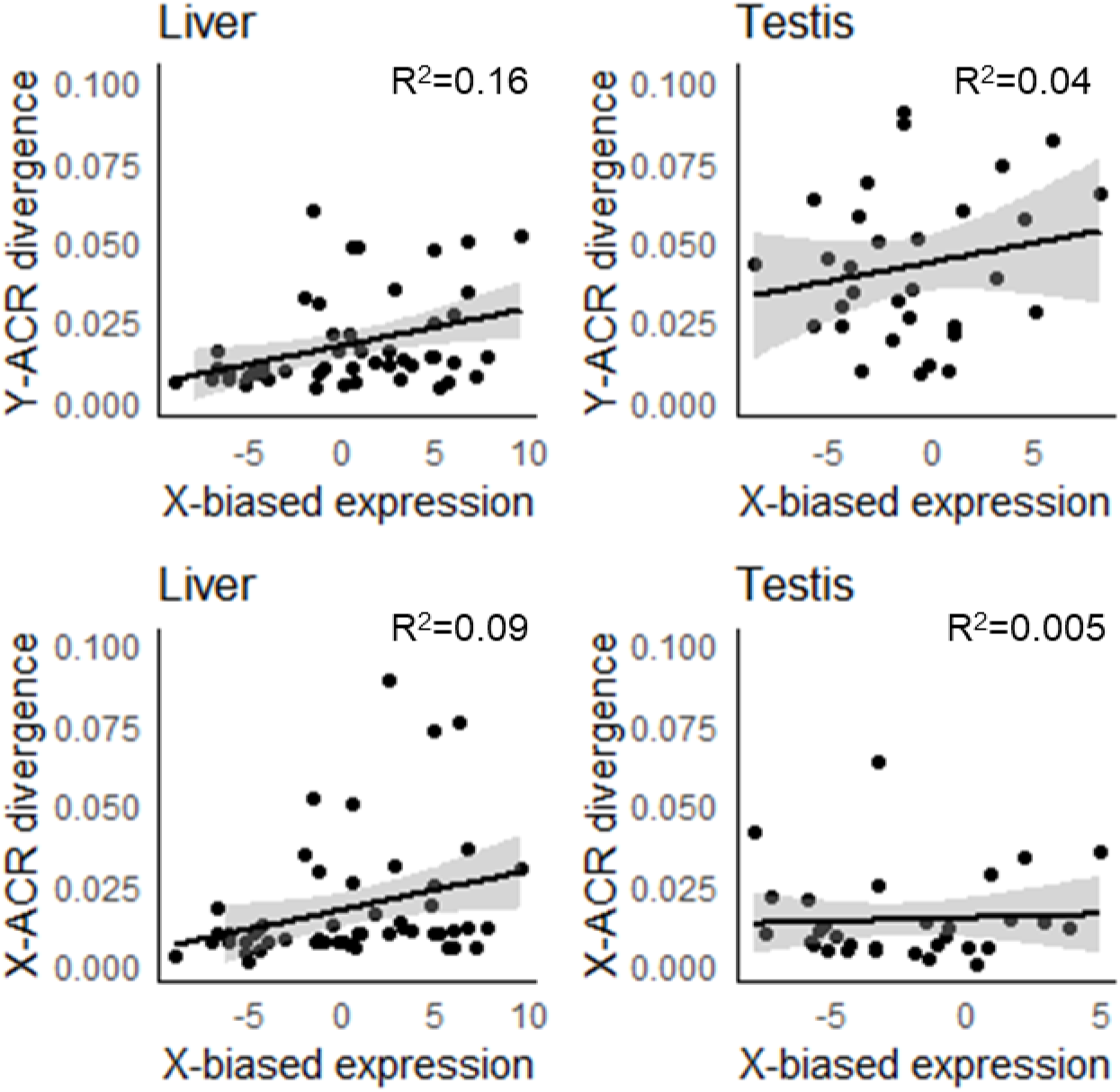
Gametolog expression level compared to accessible chromatin region divergence. ACR divergence on the X and Y chromosomes is correlated with X-biased gene expression in both liver and testis tissues. RNA-seq transcript counts that uniquely mapped to the X and Y chromosome were quantified to determine allele-specific expression for all gametologs. Gametolog expression was compared to the average divergence of Y-specific (A and B) and X-specific (C and D) mutations for ACRs within 50 kb of expressed genes. (A). Y liver: R= 0.40, P = 0.004, N = 51; (B.) Y Testis: R=0.201, P = 0.26, N = 31; (C). X Liver: R = 0.30, P = 0.033, N = 51; (D). X Testis: R = 0.07, P = 0.94, N = 31. Shaded regions represent 95% confidence intervals based on bootstrapping. The regressions were not significantly different between the X and Y chromosome for liver (P = 0.590, Fisher transformation) or testis (P = 0.077, Fisher transformation).

A similar trend was observed in testis tissue for Y-linked ACR divergence (Figure 4), but the correlations were not significant (Figure 4B; N = 31; P = 0.26; R=0.201; Spearman’s rank correlation; 95% Confidence Interval via bootstrapping (-0.0095, 0.5706)). There are fewer testis-expressed genes in this analysis. Therefore, the lack of significance could be due to sample size. To identify if sample size had an effect, we performed a randomized down-sampling for the liver-expressed genes. Even with down-sampling, 68.0% of the permutations still produced significant correlations between ACR divergence and X-biased expression in the liver (P < 0.05; 10,000 permutations). This indicates the lack of correlation observed with the testis expressed genes is likely not due to a smaller sample size. Despite having greater ACR divergence in testis compared to liver, 22 gametologs exhibited Y-biased expression in testis (Table 3). This suggests Y ACR evolution also contributes to up-regulation of male-specific genes.

**Table 3.**
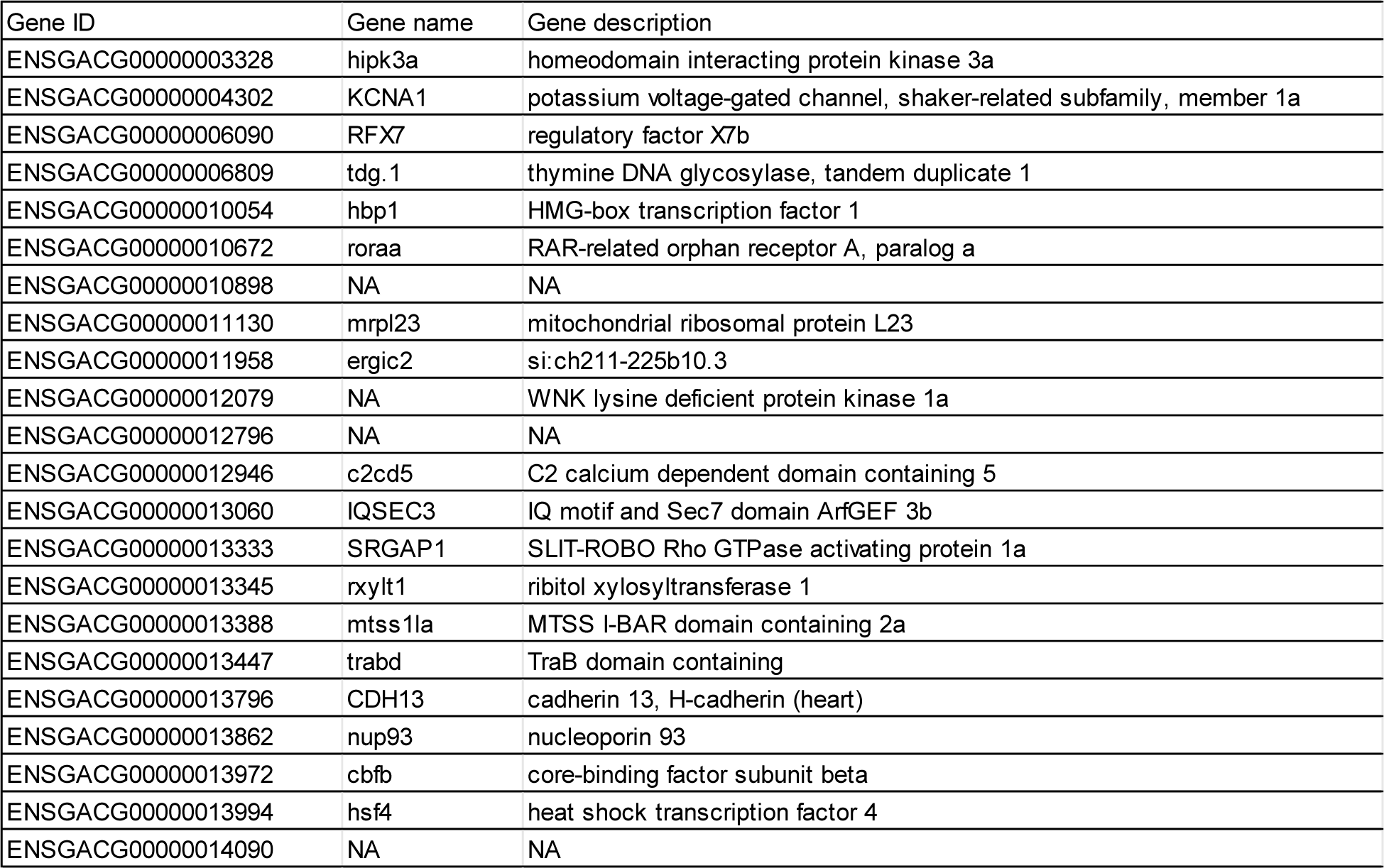
Gametologs with Y-biased expression in testes.

### *Cis*-regulatory evolution is not correlated with deleterious mutations in coding regions

The downregulation of Y-linked gametologs could be adaptive if loss of expression follows the accumulation of deleterious coding substitutions (Orr and Kim 1998). We searched for evidence of adaptive silencing on the Y chromosome by comparing the ACR divergence to the ratio of non-synonymous to synonymous substitutions (*d*_N_/*d*_S_) within the coding region of each gene. An adaptive silencing model would be supported if Y-linked gametologs with an elevated *d*_N_/*d*_S_ ratio also have elevated *cis*-regulatory divergence. Among genes shared between the X and Y chromosomes, we found no correlation between ACR nucleotide divergence and *d*_N_/*d*_S_ (Supplemental Figure 2; each stratum on X and Y: P > 0.05; Spearman’s rank correlation), indicating coding regions with elevated ACR divergence are not more likely to have elevated substitutions in the amino acid sequence.

Elevation of *d*_N_/*d*_S_ on the Y chromosome could indicate relaxed purifying selection, or positive selection. Combining genes that are evolving under both could mask signatures of adaptive silencing. To account for this, we split gametologs on the Y chromosome into classes based on the signatures of selection from the MK test. We grouped gametologs into categories of purifying selection (NI_Haldane_ > 0.25; N = 262), positive selection (NI_Haldane_ < -0.25; N = 41), and genes evolving through relaxed purifying selection (-0.25 >NI_Haldane_ > 0.25; N = 219). If selection was acting on ACRs to downregulate genes with deleterious mutations, we would expect to see a positive correlation between ACR divergence and *d*_N_/*d*_S_ for genes with signatures of relaxed purifying selection. In contrast to this prediction, we found a weak negative correlation between ACR divergence and *d*_N_/*d*_S_ for expressed genes under relaxed purifying selection (Supplemental Figure 3; R= -0.293; P =0.0155; N=35; Spearman’s rank correlation). These genes also did not have more X-biased expression than other genes (Supplemental Figure 4; P > 0.05; Kruskal-Wallis test). Interestingly, we did find a positive correlation between ACR divergence and *d*_N_/*d*_S_ for genes evolving under positive selection (Supplemental Figure 4; R=0.92; P=0.002; N=10; Spearman’s rank correlation). Overall, most genes on the Y seem to be evolving independently from neighboring regulatory regions, except for a small set of genes under positive selection.

We also examined whether ACR divergence was higher within genes that contained frameshift mutations, nonsense mutations, or in-frame deletions. These types of mutations are more likely to produce a non-functional or sub-optimal peptide. We found no significant difference in the number of ACR substitutions from liver and testis tissue for coding regions that contained frameshift, nonsense mutations, or deletions, compared to coding regions that did not have these putatively deleterious mutations (functional) (Supplemental Figure 5) (all pairwise comparisons P > 0.05; Mann-Whitney U test). Overall, we found no evidence of *cis*-regulatory divergence associated with deleterious coding sequence divergence, suggesting the adaptive silencing model is unlikely the major driver of *cis*-regulatory evolution on the threespine stickleback sex chromosomes.

## Discussion

The evolution of gene regulation on Y chromosomes has largely been studied in the context of RNA expression levels of gametologs (Muyle, et al. 2012; White, et al. 2015; Beaudry, et al. 2017; Muyle, et al. 2018; Martin, et al. 2019; Veltsos, et al. 2019). Over time, gametolog expression is generally lost across most genes on the Y chromosome. With the growing number of Y chromosome reference assembles and sequencing methods to quickly profile functional *cis*-regulatory regions across the genome, it has become feasible to explore the molecular evolution of *cis-*regulatory regions that have led to altered expression patterns. Here, we observed elevated nucleotide substitutions and reduced polymorphisms within *cis*-regulatory elements across the threespine stickleback Y chromosome, consistent with positive selection driving this process. We found that increased rates of *cis*-regulatory divergence were associated with X-biased expression of gametologs, suggesting loss of expression from the Y chromosome. Without liver and testis gene expression from the ninespine stickleback as an ancestral comparison, we were unable to confirm that X-biased gametolog expression is a result of downregulation of the Y chromosome or upregulation of the X chromosome. However, previous sequencing of a brain transcriptome showed expression patterns consistent with loss of Y expression rather than gain of X expression (White, et al. 2015). It is therefore likely that the X-biased expression we observed in liver and testis tissue also occurs through the loss of Y chromosome expression, similar to what has been observed in other species with degenerating Y chromosomes (Muyle, et al. 2012; Beaudry, et al. 2017; Wei and Bachtrog 2019).

Additional work will be necessary to identify the functional variants responsible for the X-biased expression pattern. We currently cannot determine whether the X-biased expression pattern is due to the accumulation of deleterious mutations from inefficient selection, adaptive substitutions that are accumulating in the ACRs, or a combination of the two.

Several theoretical models of sex chromosome evolution have been developed that predict how gene expression evolves over time. Like coding regions, *cis*-regulatory regions will accumulate deleterious mutations because of a reduced efficacy of purifying selection (Charlesworth and Charlesworth 2000; Bachtrog 2008). Without the removal of deleterious mutations, substitution rates within regulatory and coding regions approach the rates observed among non-functional intergenic sites. Theory predicts that substitution rates within regulatory regions can be elevated further through positive selection favoring beneficial mutations. The adaptive silencing model (Orr and Kim 1998) posits that Y gametologs that have accumulated maladaptive coding substitutions would be favored to be downregulated through *cis-*regulatory mutations. Only the functional X-linked gametolog would be expressed in males. An alternative model suggests that deleterious mutations accumulate in *cis-*regulatory elements throughout the non-recombining region, initially downregulating all Y-linked gametologs (i.e. degeneration by regulatory evolution (Lenormand, et al. 2020; Lenormand and Roze 2022). As deleterious mutations simultaneously accumulate within coding regions, a positive-feedback loop ensues, leading to a higher substitution rate within *cis-*regulatory elements to selectively downregulate increasingly maladaptive coding regions. Our survey of ACR evolution in threespine stickleback fish allowed us to compare with each of the previously discussed theories.

Evidence for adaptive silencing (Orr and Kim 1998) would be present if alleles with elevated rates of coding divergence between the X and Y chromosomes, were more likely silenced on the Y chromosome. Our results did not support this model. We found that the accumulation of X or Y ACR divergence had no association with relaxed purifying selection in nearby coding regions. Our results complement previous findings that have compared coding sequence evolution to gametolog expression. The level of expression from the Y chromosome is often not correlated with the overall number of deleterious mutations within coding regions (Bachtrog 2006; Bachtrog, et al. 2008; Beaudry, et al. 2017). While an elevation of *d*_N_/*d*_S_ has been previously used as a proxy for the accumulation of deleterious variants on the Y, this association could also be conflated by positive selection. Here, we showed the lack of an association still held when only looking at genes evolving under relaxed selection. Additionally, we found no correlation between divergence within an ACR and the functional state of a coding region (i.e. containing frameshift or nonsense mutations). We found *cis*-regulatory evolution was correlated with expression differences, but that ACR divergence accumulated independent of coding divergence. A correlation between nucleotide substitutions in *cis*-regulatory regions and deleterious mutations in coding regions may be observed if ACRs and gene expression are profiled from additional tissues. In addition, we assigned ACRs to coding sequences based on overall proximity to a gene. Although this method of annotating ACRs is commonly used to assign *cis*-regulatory regions to genes (Connelly, et al. 2014; Alexandre, et al. 2018; Ricci, et al. 2019), it is possible that some ACRs interact with other genes through long range interactions based on chromatin configuration (Mifsud, et al. 2015; Schoenfelder and Fraser 2019), reducing our ability to detect an association between deleterious mutations in coding regions and nucleotide substitutions in ACRs. Also, the number of substitutions within an ACR may not be a fully accurate predictor of the overall ability to silence a given gametolog on the Y chromosome. If only a small number of substitutions are needed to ablate transcription factor binding within an ACR, we would not expect to see a strong correlation between the number of deleterious mutations within coding sequences and the number of substitutions within ACRs. Additional functional work will be necessary to explore these alternatives.

Elevated substitution rates in *cis*-regulatory regions quickly following recombination suppression would be consistent with the model of degeneration by regulatory evolution (Lenormand, et al. 2020; Lenormand and Roze 2022). Silencing of Y-linked alleles occurs first in this model, followed by the accumulation of deleterious mutations within coding regions. We found that putative *cis*-regulatory elements exhibited signatures of positive selection on the Y chromosome across all three evolutionary strata, including the two youngest strata that still contain most of the ancestral gene content (Peichel, et al. 2020). The degeneration by regulatory evolution model predicts that the substitution rate within *cis-* regulatory elements will accelerate once deleterious mutations begin to accumulate within coding regions. This will further suppress gametolog expression from the Y chromosome, masking maladaptive coding mutations. Consistent with this model, our results show *cis-*regulatory regions have stronger signatures of positive selection compared to autosomal ACRs, as well as both coding and non-coding regions on the sex chromosomes. This suggests there is selection to downregulate Y-linked alleles in liver. Alternatively, substitutions in ACRs could reflect adaptation to upregulate genes on the Y chromosome, to maintain expression or to gain male beneficial expression (Skaletsky, et al. 2003; Martínez-Pacheco, et al. 2020; Shaw and White 2022). Non-adaptive explanations should also be carefully considered as alternatives to adaptive models. For example, elevated substitution rates could be observed in Y-linked ACRs if they have higher mutation rates compared to synonymous and silent intergenic sites. Indeed, it has been suggested that male-biased mutation rates, due to increased cell division (Link, et al. 2017), or specialized DNA repair pathways (Chang, et al. 2022), could shape the evolution of the Y chromosome. These mutations could fix at a faster rate within populations and produce false adaptive signatures because of the smaller effective population size of the Y chromosome. However, the threespine stickleback fish Y chromosome does not show evidence of male-biased mutation rate at synonymous sites (White, et al. 2015), and thus it remains unclear how these mutation biases would increase substitution rates uniquely at accessible chromatin regions.

The degeneration by regulatory evolution model also indicates dosage compensation should evolve simultaneously for genes that are under strong stabilizing selection to maintain expression levels (Lenormand, et al. 2020). Degeneration of *cis*-regulatory elements on the Y chromosome and loss of expression should select for upregulation of the gametolog on the X chromosome to compensate for dosage loss. In this scenario, nucleotide substitutions should accumulate within ACRs on the X chromosome. We found that X-linked ACRs had a depletion of polymorphism, consistent with positive selection. Unlike Y-linked ACRs, the X-linked ACRs had low divergence over longer evolutionary timespans between ninespine and threespine stickleback fish. This suggests that selection has acted on some X-linked sites more recently within threespine stickleback populations to fix beneficial regulatory mutations. This distinction is consistent with Y-linked ACR divergence accumulating first, as predicted by the degeneration by regulatory evolution model (Lenormand et al. 2020). Selection on X-linked *cis*-regulatory elements within ACRs may still be ongoing. A similar finding was observed on the neo-sex chromosomes of *Drosophila miranda*, where the neo-X chromosome was enriched for signatures of selective sweeps, while the ancestral X chromosome region was not (Bachtrog, et al. 2009).

We found the strongest signatures of positive selection on the X chromosome were from testis ACRs. Testis ACRs may have faster rates of sequence divergence on both the X and Y chromosome, from processes similar to the faster X-effect (Charlesworth, et al. 1987; Vicoso and Charlesworth 2009). There is widespread evidence for accelerated rates of coding evolution on X chromosomes compared to autosomes, especially in male specific tissues (Baines, et al. 2008; Meisel and Connallon 2013; Parsch and Ellegren 2013; Larson, et al. 2016; Kopania, et al. 2022), as the X-linked mutations in hemizygous regions are always exposed to selection in males. Expression of X-linked gametologs may also evolve at faster rates (Khaitovich, et al. 2005; Brawand, et al. 2011; Meisel, et al. 2012b; Coolon, et al. 2015; Kopania, et al. 2022), presumably due to mutations in *cis*-regulatory regions. The elevated substitution rate we observed on the X chromosome in testis ACRs may reflect male-beneficial selection to alter expression levels that is not necessarily connected to dosage compensation.

While we observed molecular signatures of positive selection on ACRs, we did not find sufficient evidence to suggest selection is associated with elevated X expression, as predicted by the degeneration by regulatory evolution model. While selection for dosage compensation could be ongoing, previous findings suggest that most genes on the threespine stickleback Y chromosome can be lost without consequence. Within the oldest stratum, over half of the Y-linked gametologs have been completely lost and males exhibit half expression, relative to females (White, et al. 2015). Even across species with chromosome-wide dosage compensation mechanisms, many genes are not compensated completely (Cotton, et al. 2013; Tukiainen, et al. 2017), which suggests that loss of Y expression is not always deleterious. The dosage sensitivity of each sex-linked gene may be essential for predicting how rapidly *cis*-regulatory regions evolve on both sex chromosomes. In the degeneration by regulatory evolution model, if stabilizing selection is relaxed, chromosome-wide degeneration in *cis*-regulatory regions still occurs, the genes just do not evolve dosage compensation (Lenormand, et al. 2020). Our results provide empirical support that *cis*-regulatory degeneration can occur without dosage compensation.

Expression balance of dosage sensitive genes can also be maintained if the Y-linked gametolog does not degenerate. The coding sequence of haploinsufficient genes on Y chromosomes have been shown to be maintained through purifying selection. These genes presumably cannot be lost from the Y chromosome (Bellott, et al. 2014; Bellott, et al. 2017; Peichel, et al. 2020; Bellott and Page 2021). For these genes, if Y-linked *cis-*regulatory elements accumulated deleterious mutations, ancestral expression patterns could be maintained if compensatory transcription binding sites evolved. This would also result in signatures of increased substitution rates with ACRs on the Y chromosome, relative to intergenic regions. This type of compensatory evolution has been proposed for functionally critical genes in mammals (Chaix, et al. 2008; Vermunt, et al. 2016), and *Drosophila* (Landry, et al. 2005; Arnold, et al. 2014; Signor and Nuzhdin 2018). Functional analysis of what transcription binding sites are affected by Y-linked mutations could help identify which mutations are leading to down-regulation of gametologs compared to those that maintain ancestral expression.

Recent models indicate inversions that suppress recombination between sex chromosomes can be selected to be retained within populations if sex-specific *trans*-acting regulators evolve to maintain optimal expression in both sexes (i.e. dosage compensation) (Lenormand and Roze 2022). This is based on the idea that if divergent X- and Y-linked *cis*-regulators were to recombine, it would lead to maladaptive expression levels in males or females. The evolutionary strata on the threespine stickleback Y chromosome evolved from at least three nested inversions that suppressed recombination (Peichel, et al. 2020). Unlike this recent model, our results suggest these inversions were likely not under strong selection to be retained initially because of early evolution of dosage compensation. Chromosome-wide dosage compensation does not occur in the threespine stickleback (White, et al. 2015) and we did not observe a corresponding evolution of X-linked upregulation. Interestingly, the excess of X-linked mutations observed in testis ACRs, may be reflective of a greater abundance of rapidly evolving sex-specific *trans*-regulators in the testes. However, additional work will be needed to identify the role that autosomal *trans*-regulators play in modifying gene expression of sex chromosomes in each sex, to test other aspects of the degeneration by regulatory evolution model.

Although we focused on the degeneration of *cis-*regulatory elements on the Y chromosome, it is important to note that Y chromosomes become masculinized over time and *cis-*regulatory elements may also be under positive selection for male-specific neo- or sub-functionalization (Soh, et al. 2014; Mahajan and Bachtrog 2017; Bachtrog, et al. 2019; Martínez-Pacheco, et al. 2020; Peichel, et al. 2020; Chang, et al. 2022). Some of the ACRs we found with high divergence may actually reflect functional *cis*-regulatory regions associated with testis-specific genes. Interestingly, we found signatures of positive selection in ACRs near testis-expressed genes. Some of these genes were enriched for Y-biased expression. These findings provide evidence that ancestral gametolog expression levels can also change presumably through gain of new testis-related functions. Additional work will be necessary to identify the novel function these regulatory regions provide during spermatogenesis. An increase in testis expression was also found for ancestral gametologs in a comparative study across 17 species of mammals (Martínez-Pacheco, et al. 2020). In these species, most Y-linked gametologs gained novel testis expression patterns, compared to their X-linked alleles. Rapid sex-linked *cis*-regulatory evolution may therefore be a universal phenomenon across species.

Y chromosomes often have much lower diversity overall, relative to neutral expectations (Lawson Handley, et al. 2006; Wilson Sayres, et al. 2014). This pattern has been attributed to purifying selection removing deleterious mutations and the linked neutral variation throughout the non-recombining region (Wilson Sayres, et al. 2014; Wilson Sayres 2018). Our findings revealed that nucleotide diversity on the threespine stickleback Y chromosome is also much lower than neutral expectations. Our results suggest positive selection within ACRs could also be an important driver of reducing nucleotide diversity on the Y chromosome within populations. One important consideration is whether sex-biased demography may affect Y-linked diversity. If males are more variable in reproductive success than females, this could lower the expected effective population size of the Y chromosome, reducing nucleotide diversity. Little is known about the operating sex-ratios of stickleback fish. Some populations exhibit sex ratios that are female biased (Rollins, et al. 2017). This could contribute to the low nucleotide diversity we observed. However, simulations in human populations revealed that even drastic shifts in sex ratio could not entirely explain the low diversity observed on the Y chromosome (Wilson Sayres, et al. 2014).

## Conclusion

Together, our results provide evidence of positive selection driving accelerated rates of nucleotide substitution in *cis*-regulatory elements. Signatures of positive selection, even in the youngest two strata, indicate that *cis*-regulatory evolution can proceed rapidly following the suppression of recombination, leading to reduced gene expression from the Y chromosome. These results support some aspects of recent models of sex chromosome evolution, where *cis-*regulatory degeneration and silencing occurs first on the Y chromosome. However, we found this can occur in the absence of dosage compensation, contrasting other assumptions of these models. Improvements in functional annotations of regulatory regions as well as an ever-growing collection of high-quality Y and W assemblies will allow continued empirical testing of new regulatory models of sex chromosome evolution.

## Materials and Methods

### Molecular evolution of accessible chromatin regions and coding regions

We used accessible chromatin regions (ACRs) from two different tissues. Liver ACRs were previously identified from two replicates (Naftaly, et al. 2021) (NCBI BioProject PRJNA667175). We also collected testes from two juvenile males (∼4.4 cm in standard length) of laboratory-reared threespine stickleback fish, originally isolated from Lake Washington (Seattle, Washington, USA) (NCBI BioProject PRJNA686097). The testis cells were immediately dissociated through homogenization in 1X PBS containing proteinase inhibitor cocktail (PIC, cOmplete tablets Roche) (PBS+PIC). The cells were fixed with 16% formaldehyde and washed twice with PBS+PIC, followed by lysis in 1M Tris-HCl, pH=8, 0.5M EDTA, 10% NP-40, 50% glycerol/molecular grade H_2_O, and 1X PIC. Nuclei were stained with DAPI and counted with a hemocytometer. We diluted samples to 60,000 – 80,000 nuclei. ATAC-seq library preparation was conducted using previously established protocols (Lu, et al. 2017; Naftaly, et al. 2021).

The libraries were sequenced on an Illumina NextSeq (2 x 150 bp; Georgia Genomics and Bioinformatics Core). Reads were trimmed with Trimmomatic (v. 0.36) (Bolger, et al. 2014), using a sliding window of four bases, trimming the remainder of the read when the average quality within a window dropped below 20. Residual adapter sequences were removed using Trimmomatic ILLUMINACLIP. Reads were filtered for a minimum length using MINLEN:30. We aligned the trimmed reads using Bowtie2 (v. 2.4.1) with default parameters (Langmead and Salzberg 2012). We filtered for alignments with a mapping quality greater than 20 using SAMtools (v1.14). We also removed PCR duplicates by using the MarkDuplicates function in Picard (https://github.com/broadinstitute/picard).

We used a Tn5 control sample to normalize ATAC-seq reads to remove the effect of Tn5 bias. For the control, genomic DNA was extracted from a caudal fin clip of one male fish using a standard phenol-chloroform extraction. The ATAC-seq library preparation, whole genome sequencing library preparation, and sequencing (Illumina HiSeq 2 x 150 bp) of the control sample were completed by GENEWIZ (New Jersey, USA). Whole genome sequencing was conducted in order to show the DNA sample was of sufficient quality to construct the Tn5 bias control. For the whole genome sequencing, we recovered over 232 million reads with an average quality score of 38.59, indicating a high-quality sample. We trimmed residual adapters and low-quality sequences using Trimmomatic as previously described. Trimmed reads were aligned to the threespine stickleback genome using Bowtie2. The read coverage per base pair was calculated using BEDTools (v2.29, -d) (Quinlan and Hall 2010). The whole genome sequencing sample had an average read depth of 90x where only ∼10% of the genome was supported with less than ten reads. Within these regions, only 6% had zero reads per base pair, indicating 94% of the genome could be queried for biased integration of Tn5. The Tn5 bias control produced over 177 million reads. The reads were trimmed from these reads using Trimmomatic with the same parameters. The trimmed reads were aligned to the threespine stickleback genome using Bowtie2 with default parameters. In order to use previous alignment coordinates between the X and Y chromosome, we used the v. 4 genome build of the X chromosome (Peichel, et al. 2020). We used the v. 5 genome build of chromosome 18 (Nath, et al. 2021). Reads mapping to the mitochondria and unscaffolded regions were removed. PCR duplicates were removed using MarkDuplicates from Picard.

We evaluated concordance between replicates in several ways. We first searched for enrichment of ATAC-seq reads around the transcription start sites (TSSs) of expressed genes. Iso-Seq long read sequencing was previously conducted in threespine stickleback fish to curate accurate transcription start site annotations across multiple tissues (Naftaly, et al. 2021). We used deepTools (v3.5.1) computeMatrix reference-point (Ramírez, et al. 2016) on the ATAC-seq alignments to assay read depth 3kb around the complete set of annotated transcription start sites. We plotted enrichment of ATAC-seq reads around TSS and compared to RNA-seq expression using deepTools plotHeatmap with default settings, except that we used the -sortUsingsamples to sort the regions by expression. We found strong enrichment around TSSs, across both replicates for the same set of expressed genes (Supplemental Figure 6). ACRs were called for each replicate using MACS2 (v2.2.7.1) callpeak with the –keep--dup all parameter, and read depth was normalized with Tn5 control sequencing with the -c parameter (Zhang, et al. 2008). The majority of ACRs were shared across replicates (liver: 67% and testis: 61%) We then used the P-value of each MACS2 peak to calculate the Irreproducibility Discovery Rate (IDR) (https://github.com/daniel-shaw1/Regulatory_divergence_paper/blob/main/Supplemental/IDR.r), (Supplemental Figure 7). The IDR determines the probability that a peak is reproducible between replicates, due to chance (Qunhua, et al. 2011; Landt, et al. 2012). Replicates that have IDR < 0.05 are generally considered to be highly reproducible. However, the IDR test is considered very stringent (Jalili, et al. 2015), potentially removing many true regions of increased accessibility. Because both replicates were exhibited high concordance, we pooled the aligned reads from both replicates and called peaks from the joint alignment using MACS2 as previously described, as pooling raw reads has been previously reported (Reske, et al. 2020; Yan, et al. 2020) as a strategy to find high quality enriched regions. We found that most ACRs (liver: 79%, testis: 75%) from the pooled sample overlapped with low scoring ACRs from the individual replicates (IDR < 0.05). This suggested a majority of ACRs in the pooled dataset were of high quality in the individual replicates.

To define an ancestral set of ACRs, we mapped the nucleotide sequence of each non-coding ACR on the X chromosome to the homologous region on the Y chromosome using previously generated alignment coordinates (Peichel, et al. 2020). A similar approach has been used previously to define the ancestral set of coding regions, aligning X-linked gene annotations to the Y chromosome (Peichel, et al. 2020). We also attempted to identify Y-linked ACRs and calculate divergence in a similar method as we identified with X-linked ACRs. However, we found that MACS2 identified significantly fewer enriched regions of accessible chromatin on the Y chromosome compared to the X chromosome. The few ACRs that were identified also aligned to multiple regions on the X chromosome, suggesting these regions were repetitive. Using the X chromosome ACRs, we identified orthologous regions between the X chromosome, Y chromosome, and orthologous autosome (chromosome 19) from the ninespine stickleback fish *(Pungitius pungitius*; (Varadharajan, et al. 2019) using BLAST+ (blastn v. 2.11.0) with default blastn parameters and -perc_identity set to 75. (Camacho, et al. 2009). To ensure high quality alignments, we filtered for uniquely mapping alignments that also had a bit score greater than 100. We used BLAST+ alignments to extract ACRs and the proximal 50bp upstream and downstream, to create multiple sequence alignments for downstream analysis. We aligned the threespine stickleback X and Y ACR sequences to the ninespine stickleback autosome sequence using MUSCLE (v 3.8.1551) (Edgar 2004) with default parameters. We used MUSCLE to create a multiple sequence alignment between the X sequence, Y sequence, and the sequence from the orthologous autosome in order to identify X- and Y-specific variants. We called single nucleotide variants using snp-sites (v2.5.1) (Page, et al. 2016) with default parameters. We calculated the reported divergence rate by dividing the number of variants by the number of the aligned sites.

We compared ACR divergence on the sex chromosomes to ACR divergence on autosome 18, a chromosome similar in length to the X chromosome. We identified orthologous regions between the threespine stickleback autosome 18 and autosome 18 from the ninespine stickleback fish using BLAST+ (blastn v. 2.11.0) (Camacho, et al. 2009) as previously described. We called variants between autosomal regions in a similar manner as the sex chromosomes, by creating a pairwise alignment using MUSCLE (v 3.8.1551) with default parameters and called single nucleotide variants using the same python script as above. We calculated a divergence rate by dividing the number of single nucleotide variants by the size of the aligned region.

Divergence within ACRs was compared with the divergence of coding regions of neighboring genes. We identified the closest gene to each ACR by running annotatepeaks.pl in homer (v 4.11) (Heinz, et al. 2010) using default settings. Per default settings, ACRs were assigned to the closest gene within 50kb of the TSS. We verified the homer peak annotations with a custom Python script that identifies ACRs near the TSS of genes (https://github.com/daniel-shaw1/Regulatory_divergence_paper/blob/main/Supplemental/Peaksneargenes.py). Both methods were 100% concordant with each other. We further verified a subset of the annotations (200 peaks total) by visualizing the ACRs in the threespine stickleback JBrowse genome browser (https://stickleback.genetics.uga.edu/). We found that 100% of the 200 X-linked ACRs were located proximal to the predicted annotated gene (Supplemental Figure 8). For coding divergence comparisons, we used previously reported estimates of synonymous (*d*_S_) and non-synonymous (*d*_N_) divergence of gametologs between the X and Y chromosomes (Peichel et al., 2020). Coding regions with *d*_N_/*d*_S_ ratios of 99 were omitted as these represent alignments with very little sequence divergence to estimate *d*_N_/*d*_S_ accurately.

### Estimating a control substitution rate

To generate control regions, we randomly sampled non-functional intergenic regions throughout the X chromosome, Y chromosome, or chromosome 18 to estimate a neutral substitution rate. Regulatory regions tend to be GC-rich, which can be prone to higher mutation rates through mechanisms like GC-biased gene conversion or spontaneous deamination of methylated cytosines (Nesta, et al. 2021). We therefore GC-matched the randomly drawn intergenic regions with the GC content of ACRs. We calculated the GC percentage of ACRs using a modified Perl script (countbp.pl, Nicholas Navin, http://www.navinlab.com/bioperl/bioperl/gc_content.html). For each ACR, a random intergenic region was drawn from the X chromosome equal in length and with a GC content within 2.0%. Intergenic regions were defined as any region that fell outside of annotated functional regions from a combination of Ensembl annotations (Cunningham, et al. 2021), Isoseq transcripts (Naftaly, et al. 2021), Y chromosome annotations (Peichel, et al. 2020), repetitive elements (Peichel, et al. 2020; Nath, et al. 2021) and ACRs from this study. This was performed for each stratum individually using bedtools shuffle –I ACRs.bed -incl XYalignment.bed -excl annotations.bed. We extracted 10,000 sets of GC-matched intergenic regions from each of the three evolutionary strata on the threespine stickleback sex chromosomes (set size same as number of Liver ACRs: stratum one: 442, stratum two: 228, stratum three: 199; Testis ACRs: stratum one: 30, stratum two: 37, stratum three: 51). Due to a limited number of intergenic sites shared between the sex chromosomes, we sampled with replacement to align a sufficient number of intergenic regions. We identified the orthologous regions to the ninespine genome assembly using BLAST+ (blastn v2.11.0) (Camacho, et al. 2009). We filtered for alignments that mapped uniquely to chromosome 19 in the ninespine stickleback fish, had an alignment length equal to 75% of the threespine query sequence, and had a bit-score greater than 100. We generated multiple sequence alignments using MUSCLE (v 3.8.1551), for each intergenic permutation and calculated X and Y specific substitution rate as previously described.

### Estimating within population nucleotide diversity

We used whole-genome short-read sequencing of 12 males from the Lake Washington population (Washington, USA; NCBI SRA SRP137809) to estimate nucleotide diversity across the sex chromosomes. The raw reads were trimmed with Trimmomatic (v. 0.39) (Bolger, et al. 2014), using a sliding window of four bases, trimming the remainder of the read when the average quality within a window dropped below 15. The leading and trailing base pairs below quality three of every read were removed along with any residual adapter sequence. After trimming, any reads below a minimum length of 36 were discarded. We aligned the trimmed reads using Bowtie2 (v. 2.4.5) with default parameters (Langmead and Salzberg 2012). We also marked PCR duplicates by using the MarkDuplicates function in Picard (v. 2.26.10) (https://github.com/broadinstitute/picard). SNP genotyping was conducted using the GATK software package (v. 4.2.5.0). We called variants using HaplotypeCaller in GVCF mode. We then joint-called variants using GenotypeGVCFs using a genomics database created by GenomicsDBImport.

We only considered biallelic SNPs in our estimates of nucleotide diversity and we filtered for high quality genotypes using several different methods. On the sex chromosomes, sites should only have hemizygous genotypes in males in the non-crossover region outside of the pseudoautosomal region. Any sites that are heterozygous would be caused by errors in read alignment. We therefore did not consider sites that were heterozygous in the non-crossover region on the X or Y chromosomes. We also filtered out sites that exhibited too low or high of read depth, which would be indicative of alignment errors. To do this, we did not consider sites that were less than one-half or greater than double the median read depth of each chromosome (X and Y chromosomes: positions were retained if the read depth was between 3.5 and 14; chromosome 18: positions were retained if the read depth was between 6.5 and 26). Read filtering was conducted using custom Perl scripts.

Nucleotide diversity (pi) was estimated on a per-site basis (--site-pi) across each chromosome using vcftools (v. 0.1.16) (Danecek, et al. 2011)). Nucleotide diversity was averaged within 10,000 bp non-overlapping windows across chromosomes X, Y, and 18.

### Testing for positive selection using population polymorphism

We performed a modified version of the McDonald-Kreitman test (Andolfatto 2005) by treating divergence (D) and diversity (P) within ACRs as the non-synonymous factor, and non-functional intergenic sequence previously generated was used as synonymous factor. As intergenic sequence is independent of each ACR, we pooled sites from each factor as previously described (Andolfatto 2005). We counted the total number of polymorphic sites or fixed substitutions (divergence) for each factor to calculate *α*, the fraction of substitutions that are likely fixed due to positive selection. Divergence was estimated either between the threespine stickleback X and Y chromosomes or between the threespine and ninespine stickleback autosome 18. Negative alpha values were interpreted as regions that have rates of adaptive evolution near zero (Good, et al. 2013). We also used these counts to estimate the neutrality index (NI) (Rand and Kann 1996). An NI greater than one indicates an excess of ACR divergence within species, while an NI less than one indicates an excess of ACR divergence between species (or between the X and Y). An index lower than one would be consistent with positive selection. To test for significant departures from neutrality within each stratum or chromosome we performed a chi-square test to compare the rates of *d*_N_/*d*_S_ to *p*_N_*/p*_S_. Standard deviation was determined through non-parametric bootstrapping. We bootstrapped 10,000 replicates for each factor within the McDonald-Kreitman or NI test and computed both metrics for each replicate.

We also performed a McDonald-Kreitman test on coding regions. We aligned the coding sequence of all threespine stickleback X chromosome Ensembl predicted genes (build 95) to the Y chromosome and the ninespine stickleback outgroup (autosome 19; v. 7) (Varadharajan, et al. 2019) using Exonerate (v. 2.4.0) (Slater and Birney 2005) with parameters: --model coding2genome -M 2000 --bestn 1 --annotation. The Ensembl coding regions from the threespine stickleback autosome 18 were also aligned to the ninespine stickleback autosome 18. The highest scoring alignment from Exonerate was used for the MK test. Divergence and polymorphism was tallied at each position of the alignment. If a position was variable, we determined if this was a synonymous or non-synonymous mutation in the threespine stickleback by comparing to the ninespine stickleback translated codon. Sites were omitted if they contained any N characters or contained any missing genotype data among the 12 males within the VCF file. Codons were omitted if they contained any gaps from the alignment. We only considered biallelic sites when counting polymorphisms and fixed substitutions within the Lake Washington population. We used custom Perl scripts to count the total number of divergent and polymorphic sites. To compare to ACRs, we calculated *α* and NI by pooling sites across all coding regions. For gene-by-gene MK tests, we log transformed NI to include genes with a count of zero for any particular category of divergence or polymorphism and refer to the value as NIHaldane (Haldane 1956; Stoletzki and Eyre-Walker 2010).

### Comparison with allele-specific expression patterns

We measured allele-specific expression of transcripts on the X and Y chromosome using three replicate liver samples (NCBI BioProject PRJNA591630) and three replicate testis samples (NCBI BioProject PRJNA591630). We trimmed the RNA-seq reads with Trimmomatic (v. 0.36) (Bolger, et al. 2014) using a sliding window of 4 bases, trimming the remainder of the read when the average quality within a window dropped below 20 (SLIDINGWINDOW:4:20). Residual sequencing adapters were also removed using ILLUMINACLIP. We aligned RNA-seq reads using Tophat2 (v2.2.1) with default parameters (Kim, et al. 2013). We filtered for alignments with a map quality greater than 25 using SAMtools (v1.14) (Li, et al. 2009) to identify reads that map uniquely on the X and Y allele. Similar map quality filters have been used to distinguish short reads between sex chromosomes in *Drosophila* and stickleback fish (Ellison and Bachtrog 2019; Peichel, et al. 2020) and between sexes with heteromorphic sex chromosomes in guppies (Kirkpatrick, et al. 2021). Read counts were obtained using htseq-count (v 0.9.1) (Anders, et al. 2015). Default parameters were used with the addition of --stranded=no and -- nonunique all to maximize our ability to count reads on the Y chromosome. We used a custom GTF file for htseq-count that included previously determined start and end sites of all Ensembl predicted transcripts that are shared between the X and Y chromosomes (Peichel, et al. 2020). Reads were counted on the X and Y across the entire transcript as a feature. Transcripts were removed from the analysis if they had an RNA expression count of zero in all samples. For each tissue, we calculated the average read count for each gametolog across the three tissue replicates. We calculated the allele-specific expression as log_2_(X transcript average / Y transcript average).

## Data availability

The data for this manuscript is available within the short reads archive under accession numbers: PRJNA667175, PRJNA591630, PRJNA686097, SRP137809.

Scripts and bioinformatic pipelines are available at: https://github.com/daniel-shaw1/Regulatory_divergence_paper

## Acknowledgements

This work was supported by the National Science Foundation MCB 1943283 (M.A.W.), the National Institutes of Health R01GM147312 (M.A.W.), and by the University of Georgia Research Foundation (D.E.S.). We would like to thank Bob Schmitz and Casey Bergman for helpful discussions related to the project.

## Supplemental Figures

**Supplemental Figure 1.**
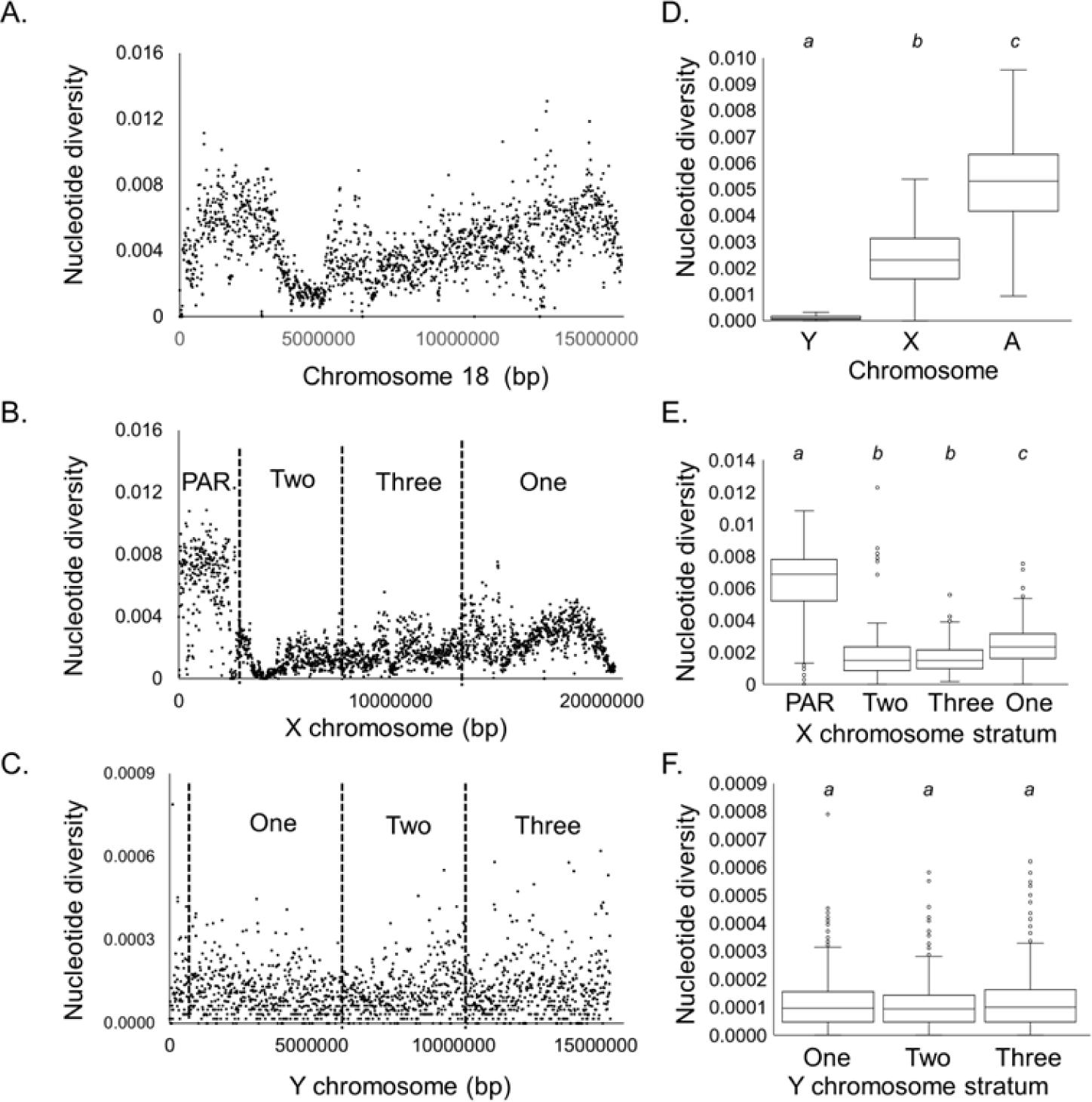
Nucleotide diversity (pi) across the sex chromosomes and representative autosome. The Y chromosome exhibits a strong reduction in nucleotide diversity consistent with a lower effective population size, and selection. Estimates of nucleotide diversity were calculated from 12 male individuals from a freshwater population. Nucleotide diversity was averaged across 10,000bp windows for A. Chromosome 18, B. X chromosome, and C. Y chromosome. D. The Y chromosome has lower diversity than both the X chromosome and chromosome 18. E. Distinct strata on the X chromosome have varying levels of the nucleotide diversity. A Kruskal Wallis test indicated differences between the groups (P < 0.001). A post-Hoc Dunn’s test and Bonferroni Correction revealed that the PAR was significantly higher from all three strata (P < 0.001). Stratum one was significantly higher than stratum two and three (P < 0.001). After removing windows that overlapped the centromeric region, stratum two and three are not significantly different (P =0.937). F. All three strata across the Y chromosome have similar estimates of nucleotide diversity (P=0.768, Kruskal Wallis test). The PAR was not analyzed for the Y chromosome as the assembly is missing most of the PAR sequence. The letters above each plot indicate the significant differences between sites within each stratum (P < 0.001). Box plots that share a letter assignment are not significantly different from one another (P > 0.001).

**Supplemental Figure 2.**
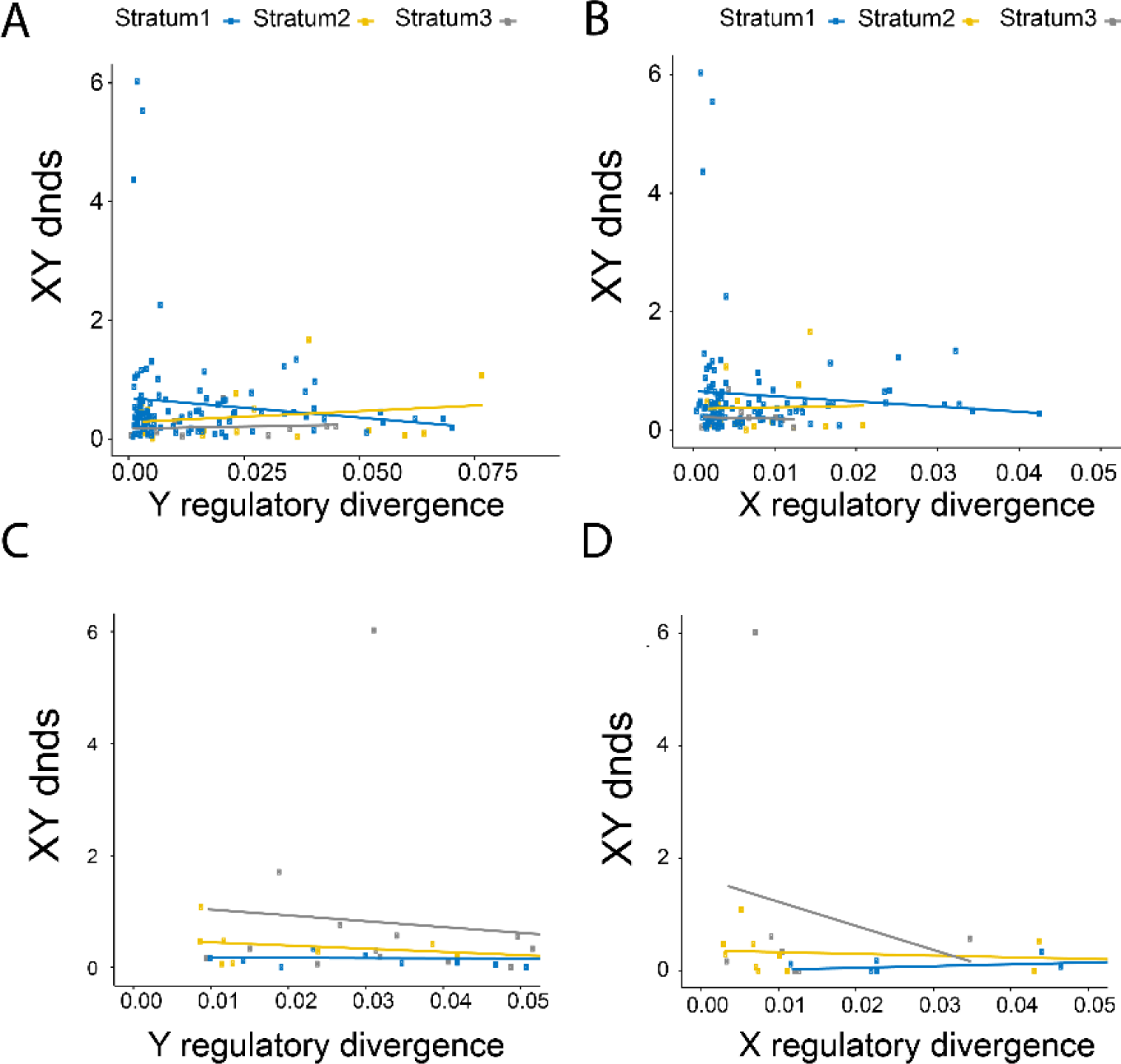
d_N_ / *d*_S_ ratios of coding regions compared to accessible chromatin region divergence. There is no correlation between *d*_N_ / *d*_S_ and ACR divergence using a Spearman’s Rank Correlation using ACRs from liver (A and B) or testis (C and D) tissue. Liver: (**A**) Y divergence: (Stratum one (Blue): R=-0.12 P = 0.230, N = 30, Stratum two (Yellow): R = 0.2, P = 0.430, N=61; Stratum three (Grey): R = 0.11, P = 0.770 N=126. (**B**) X divergence: (Stratum one (Blue): R = -0.080, P = 0.420, N=30; Stratum two (Yellow): R = 0.032, P = 0.9, N = 61; Stratum three (Grey): R = -0.054, P = 0.890, N = 126. Similar results in testis: (**C**) Y divergence: (Stratum one (Blue): R = -0.020, P = 0.650, N=14; Stratum two (Yellow): R = -0.430, P = 0.160, N = 12; Stratum three (Grey): R = -0.140, P = 0.480, N = 17. (**D**) X divergence: (Stratum one (Blue): R = 0.520, P = 0.120, N=14; Stratum two (Yellow): R = - 0.360, P = 0.210, N = 12; Stratum three (Grey): R = -0.020, P = 0.670, N = 17.

**Supplemental Figure 3.**
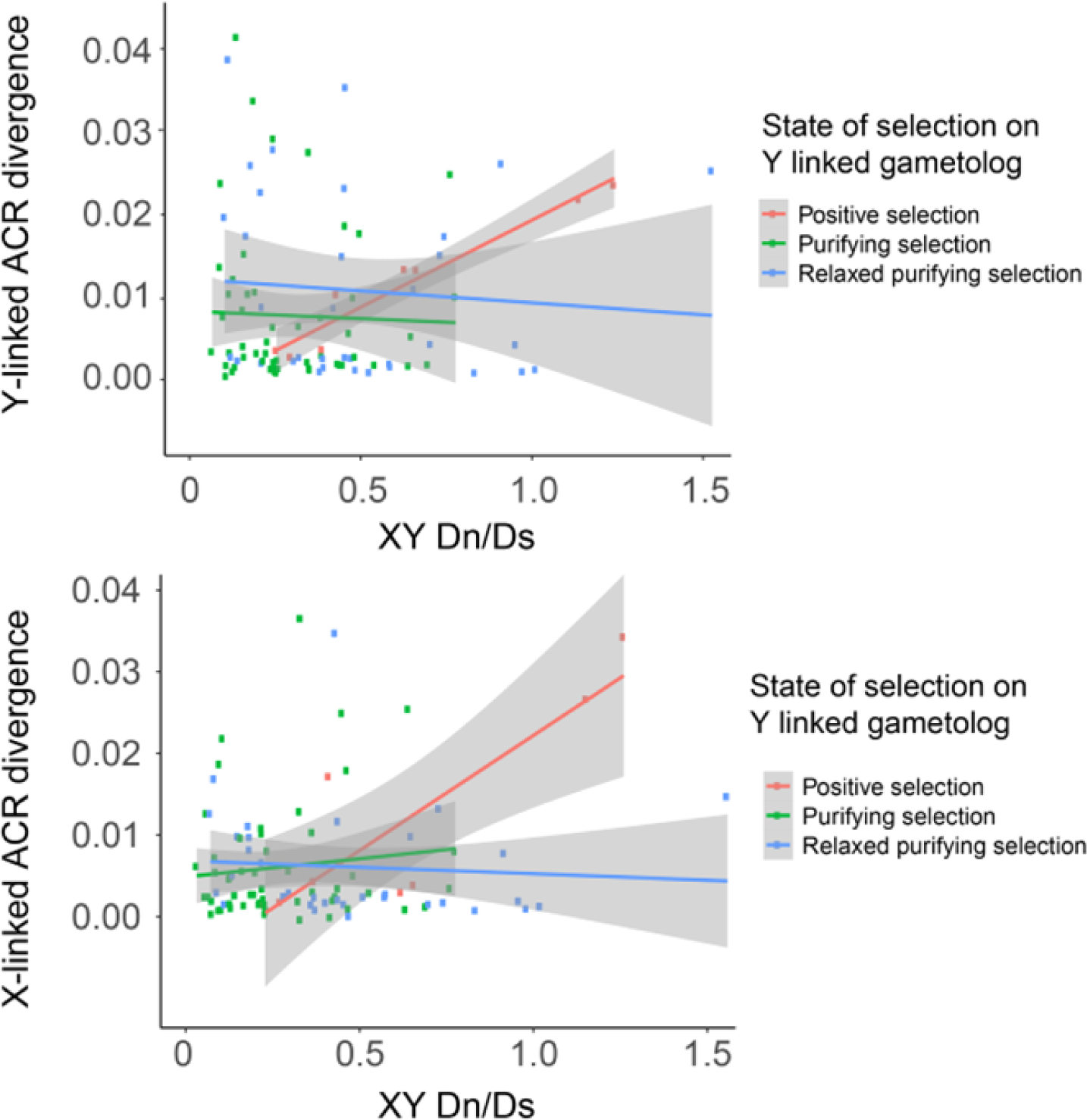
Selective constraint of genes affects correlation between ACR and proximal coding divergence. Y-linked gametologs were classified by their mode of selection based on a McDonald Kreitman Test. Genes were classified as Positive selection: NI_Haldane_ < -0.25, Relaxed purifying selection: - 0.25 <NI_Haldane_ < 0.25, and purifying selection: NI_Haldane_ > 0.25. Genes likely evolving under positive selection had Y-linked and X-linked ACR divergence significantly correlated with *d*_N_/*d*_S_ (Y chromosome: R=0.92, P=0.002, N=10; X chromosome: R = 0.81, P = 0.03, Spearman’s Rank correlation coefficient). Genes under purifying selection and relaxed purifying selection did not show correlations between ACR divergence and DN/DS (Purifying: Y chromosome: R= -0.14, P = 0.30, N=57; X chromosome: R= - 0.022, P =0.87, N=57; Relaxed purifying selection: Y chromosome: R= -0.293, P =0.0155, N=35, X chromosome: R= -0.32, P =0.063, N=35, Spearman’s Rank Correlation coefficient. Grey areas represent 95% confidence intervals. Outliers with *d*_N_/*d*_S_ > 3 were removed prior to analysis. When *d*_N_/*d*_S_ > 1 are removed the relationship for genes under positive selection is still significant (R=0.86, P=0.006).

**Supplemental figure 4.**
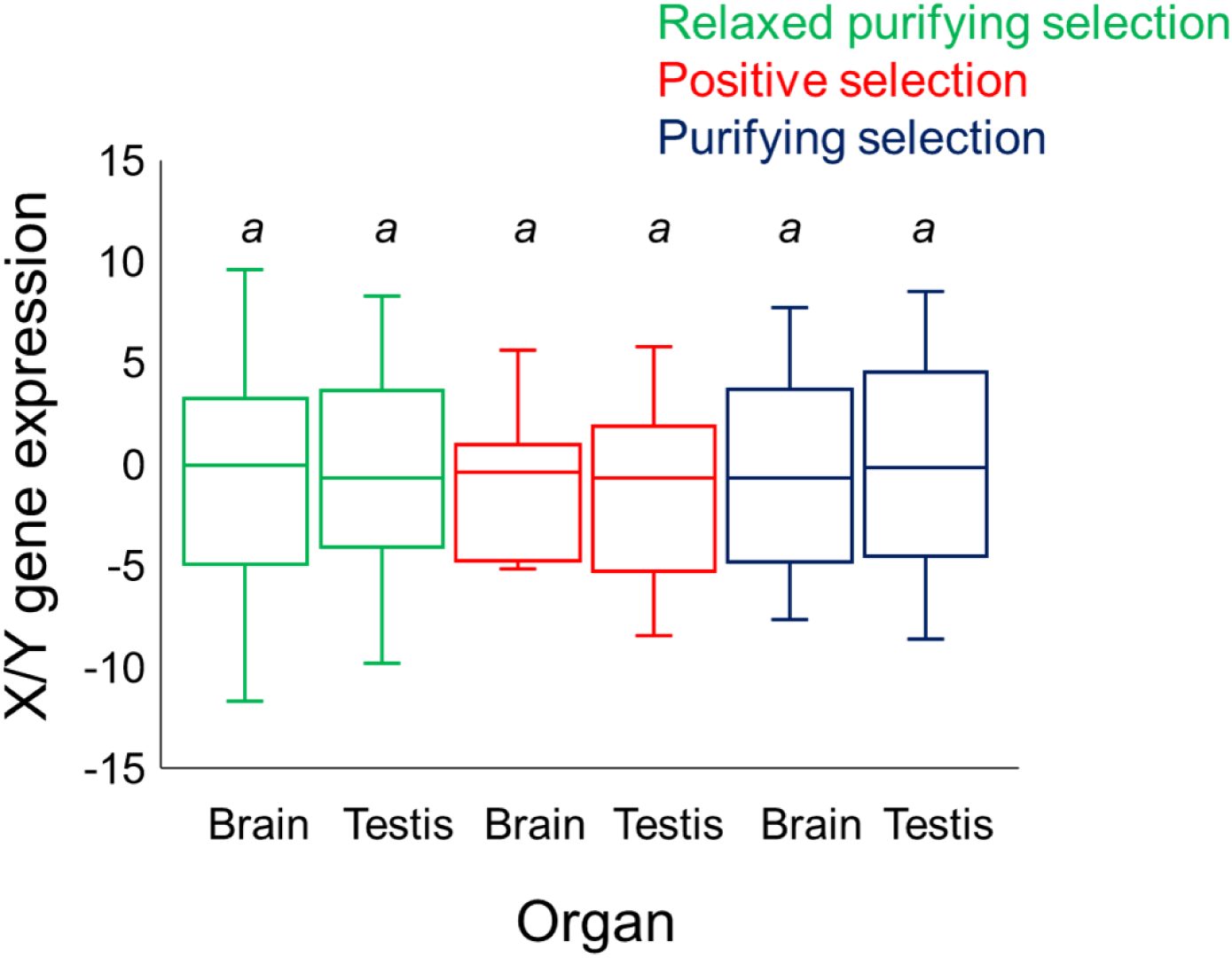
Rates of allelic expression for Y gametologs under different modes of selection. Y-linked gametologs were classified by their mode of selection based on a McDonald Kreitman Test. Genes were classified as Positive selection: NI_Haldane_ < -0.25, Relaxed constraint selection: -0.25 <NI_Haldane_ < 0.25, and purifying selection: NI_Haldane_ > 0.25. X/Y gene expression was quantified based on counts from liver and testis RNA-seq data from 3 individuals. Counts were averaged across all three replicates. Rates of X/Y expression are not significantly different for each class (Kruskal-Wallis test).

**Supplemental Figure 5.**
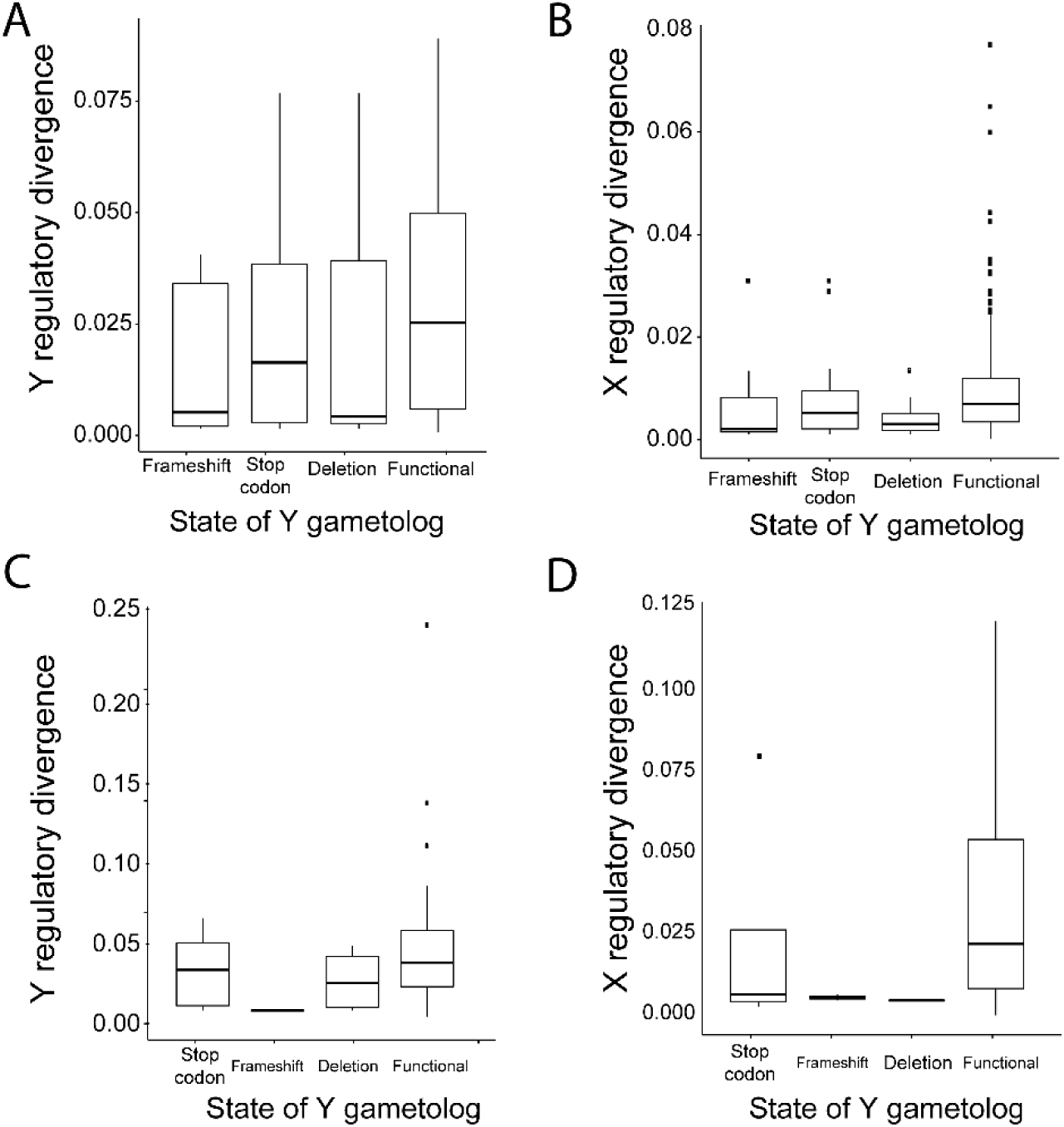
Accessible chromatin region divergence compared to functional states of Y-linked gametologs. ACR divergence was not significantly different among gametologs that contained mutations that potentially impede function (i.e., premature stop codons, frameshifts, deleted nucleotides) or are functional. Y-specific functional mutations were identified by comparing coding sequence between the Y chromosome and the outgroup autosomal coding sequence in the ninespine stickleback. ACRs were assigned to the closest gametolog within 50kb. P > 0.05 for all pairwise comparisons for both liver (A and C) and testis (B and D) for regulatory divergence on the X and Y.

**Supplemental Figure 6.**
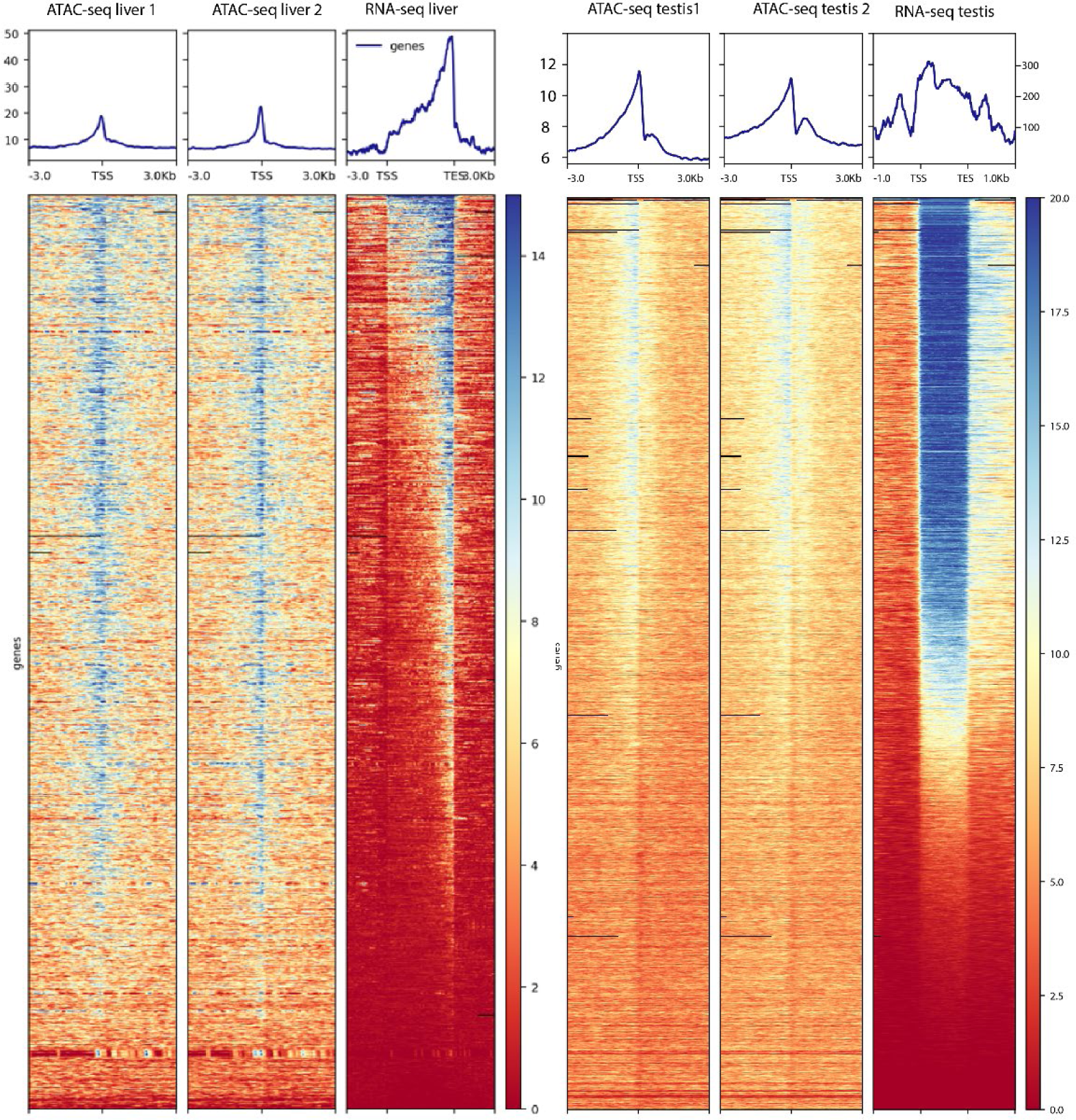
High concordance of chromatin accessibility around transcription start site of expressed genes. Transcription start sites (TSSs) for all genes were ordered by RNA-seq read depth. ATAC-seq read depth was quantified ±3 kb of TSSs. Similar ATAC-seq read depth was observed between replicates. ATAC-seq depth across replicates is a strong predictor of expression from RNA-seq from same tissue type.

**Supplemental Figure 7.**
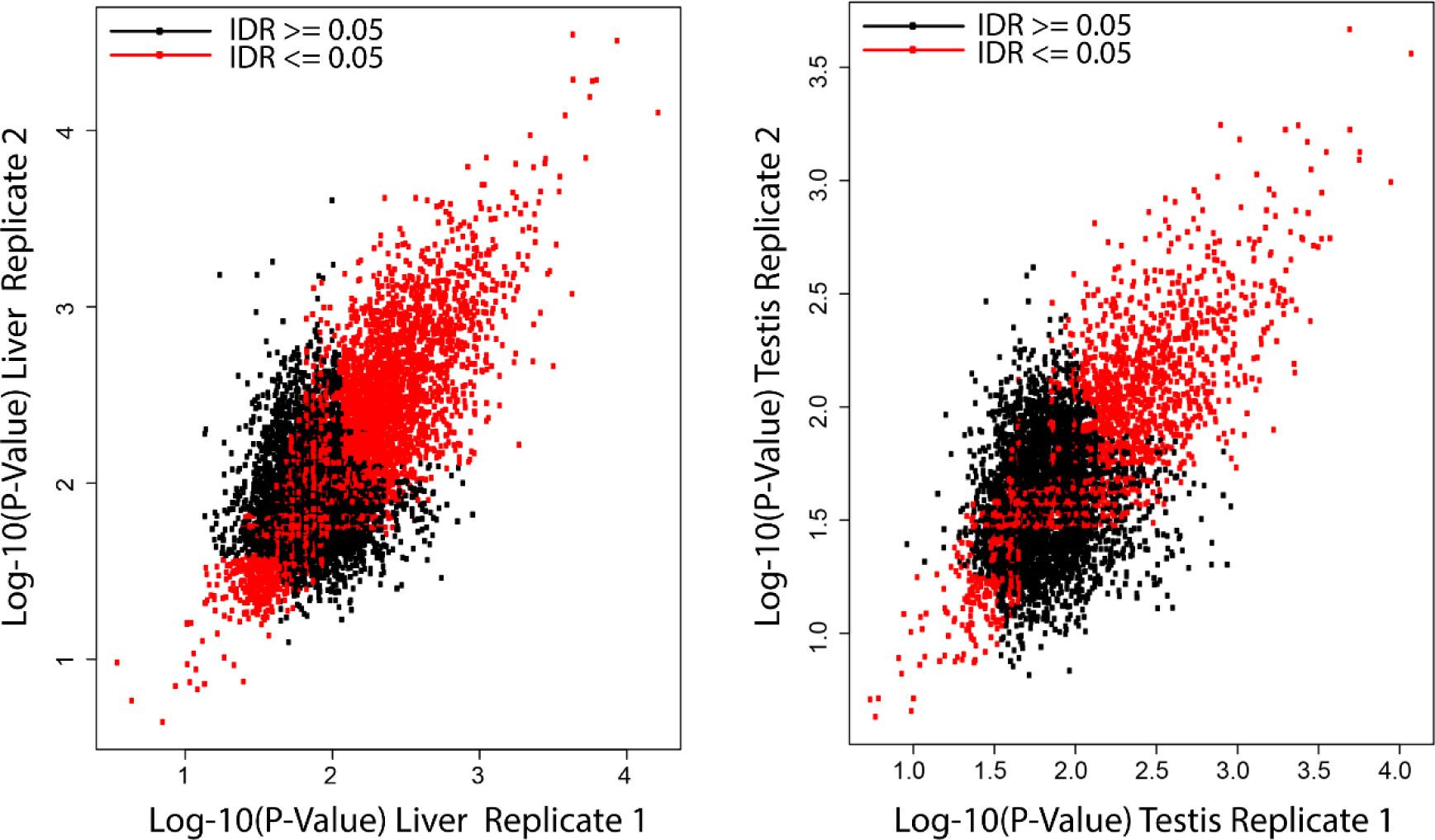
High concordance of chromatin accessibility between peaks identified in two replicates. Peaks were identified in two replicates of liver (left) and testis (right). Log_-10_(P-Value) of each shared peak is highly correlated between replicates. Irreproducible Discovery Rate was calculated by calculated the ranked significance of reproducible and irreproducible peaks between replicates. Peaks with an IDR < 0.05 are shown in red.

**Supplemental Figure 8.**
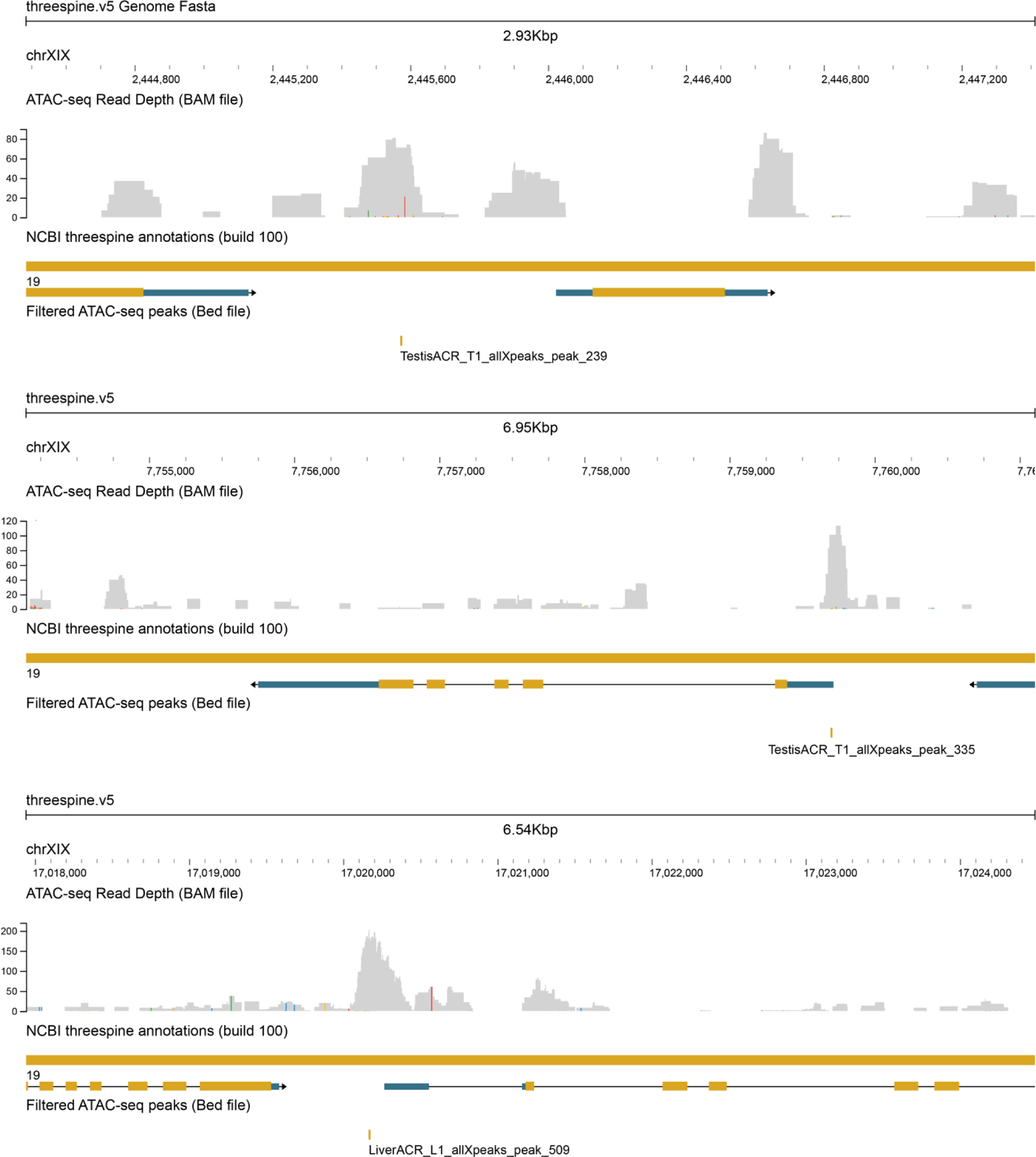
Verification of ATAC-seq proximity to annotated genes. Three representative examples of ATAC-seq read depth enriched at ACRs called with MACS2. The most proximal genes were identified with homer and with a custom python script. Annotations for a subset of 100 X chromosome ACR were visually inspected with JBrowse, to confirm that the peak was accurately assigned to nearby genes. ATAC-seq read depth is shown in grey. Bed files for the summit of each ACR identified by MACS2 is shown below the Ensembl gene annotations.

