## Supplemental material for "Positive selection drives *cis-*regulatory evolution across the threespine stickleback Y chromosome"

**Supplemental Figure 8. Verification of ATAC-seq proximity to annotated genes.** Three representative examples of ATAC-seq read depth enriched at ACRs called with MACS2. The most proximal genes were identified with homer and with a custom python script. Annotations for a subset of 100 X chromosome ACR were visually inspected with JBrowse, to confirm that the peak was accurately assigned to nearby genes. ATAC-seq read depth is shown in grey. Bed files for the summit of each ACR identified by MACS2 is shown below the Ensembl gene annotations.

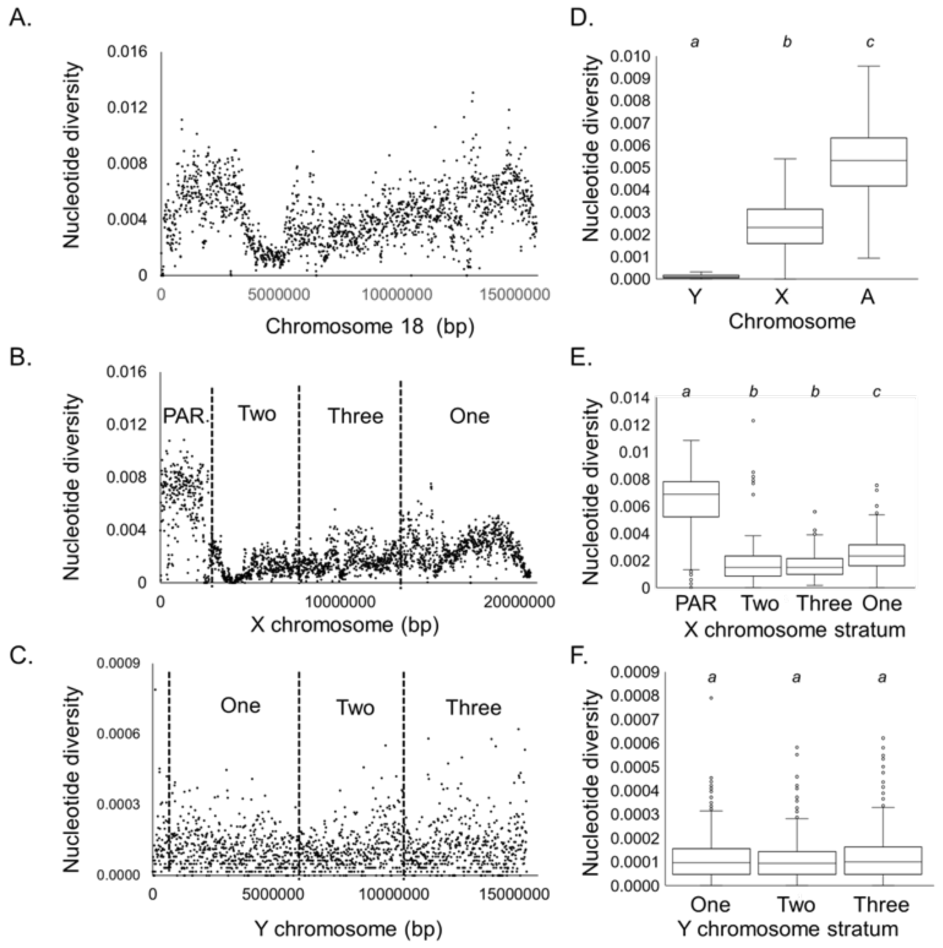


**Supplemental Figure 1. Nucleotide diversity (pi) across the sex chromosomes and representative autosome.**

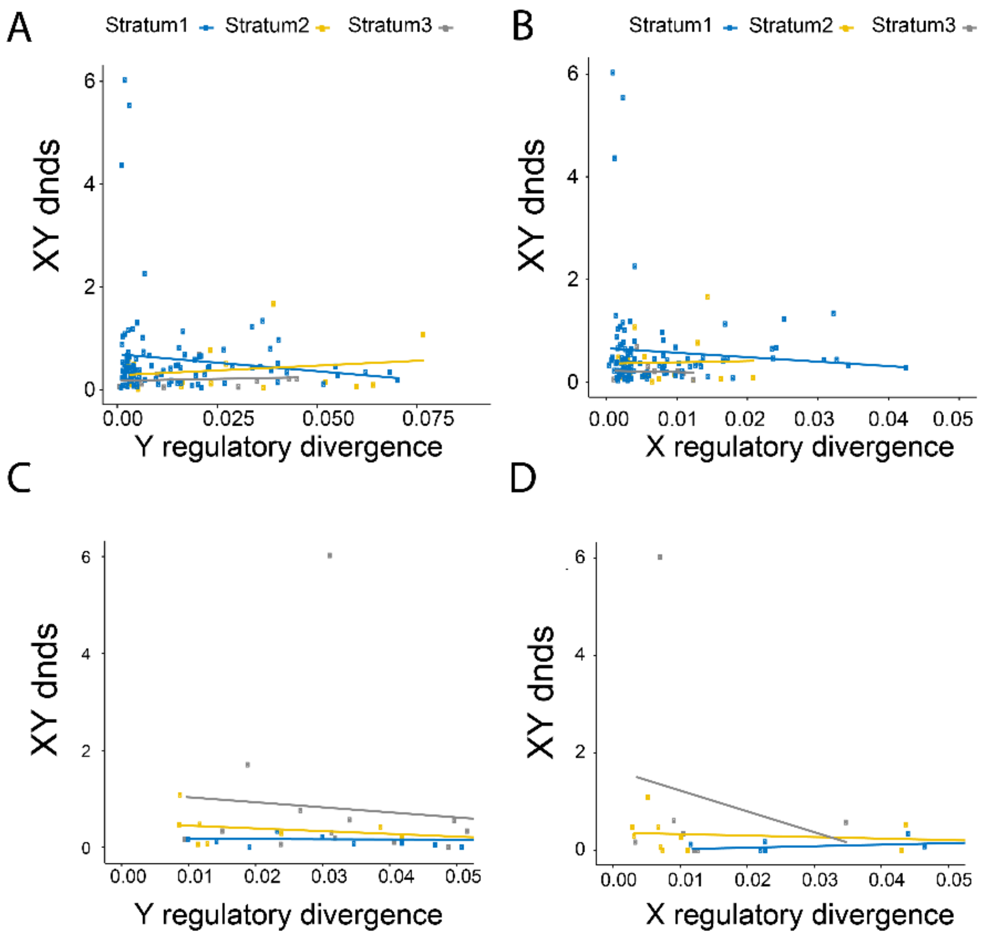


**Supplemental Figure 2.** $\boldsymbol{d_N / d_S}$ **ratios of coding regions compared to ACR divergence.**

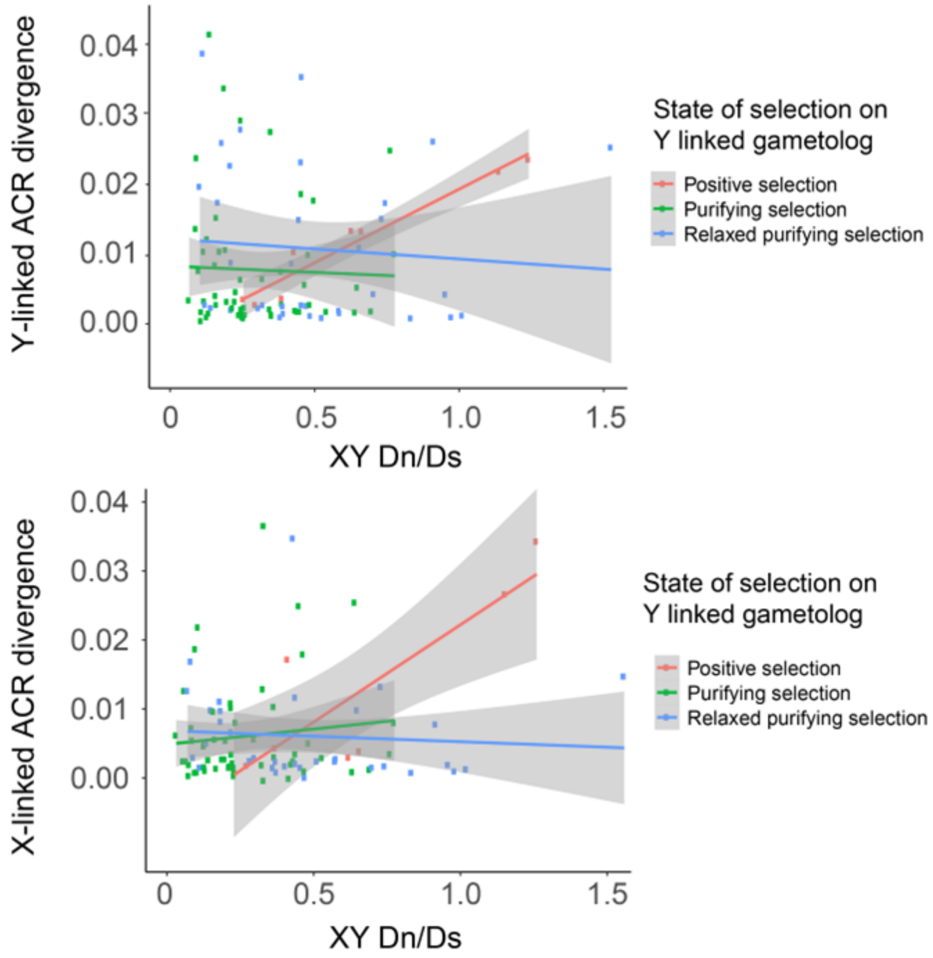


**Supplemental Figure 3. Selective constraint of genes affects correlation between ACR and proximal coding divergence.**

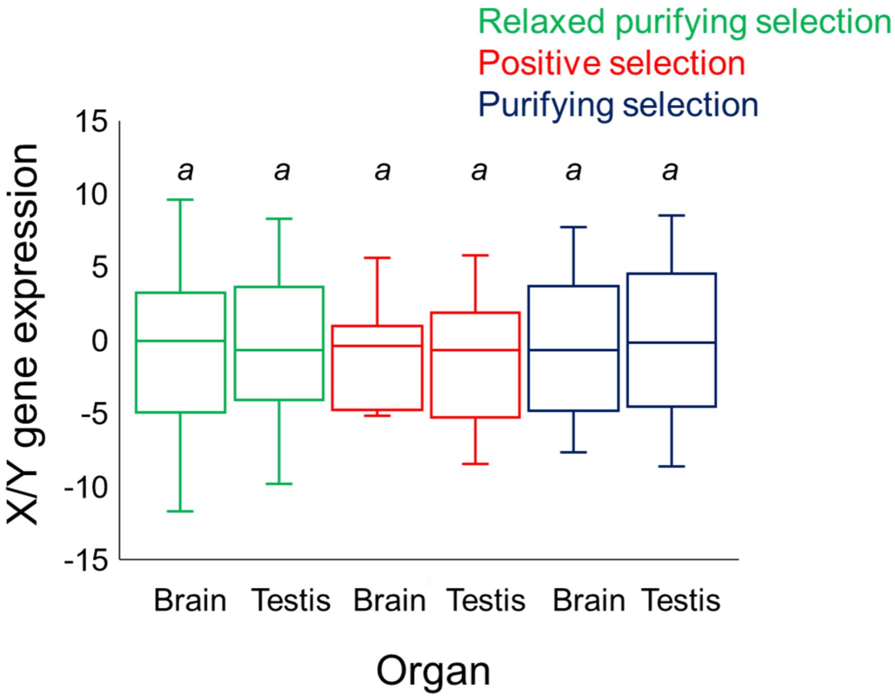


**Supplemental figure 4. Allelic-specific expression for Y gametologs under different modes of selection**

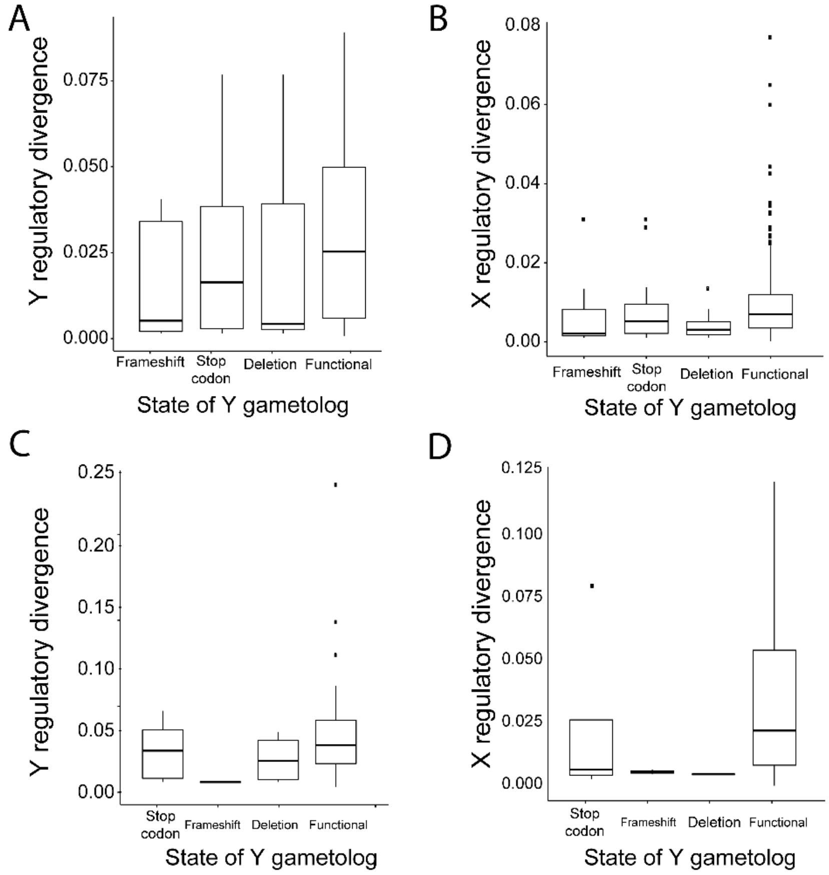

**Supplemental Figure 5. ACR divergence compared to functional states of Y-linked gametologs.**

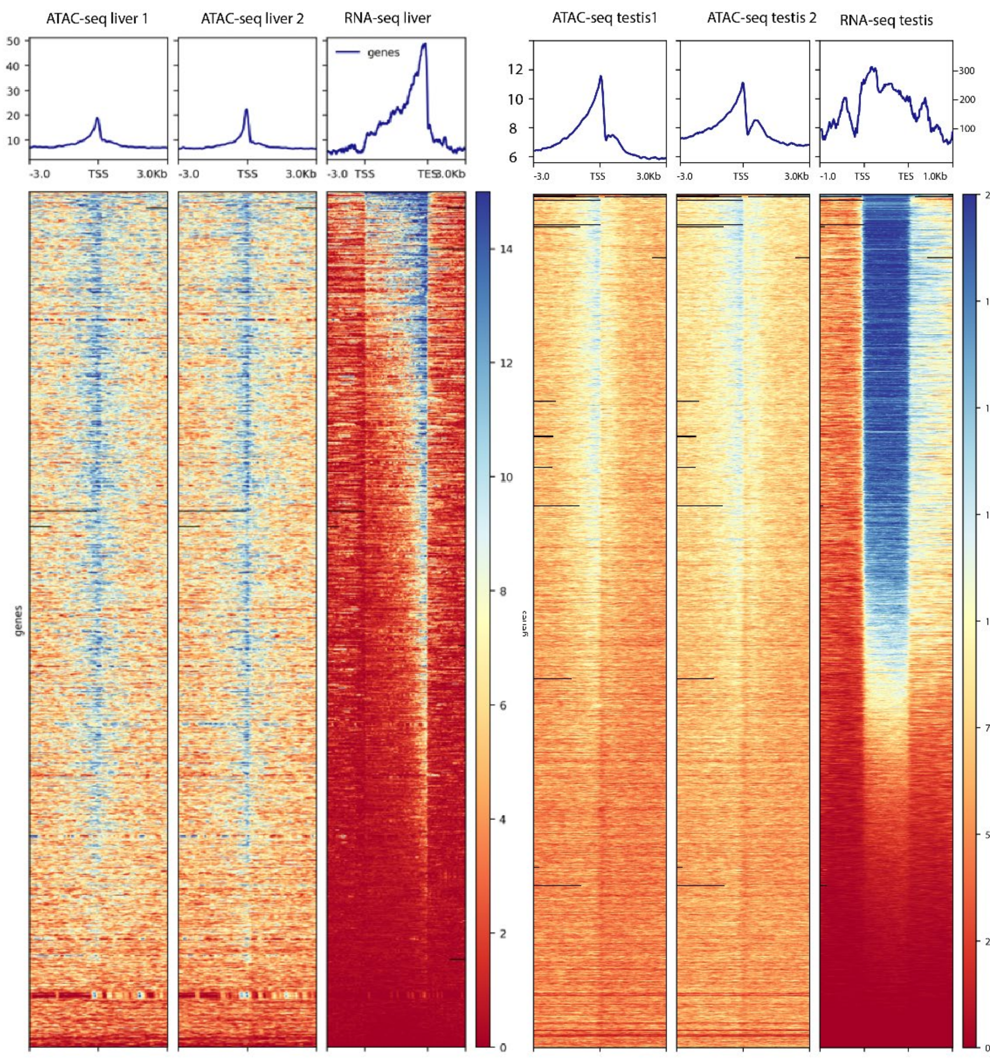


**Supplemental Figure 6. High concordance of chromatin accessibility around transcription start site of expressed genes.**

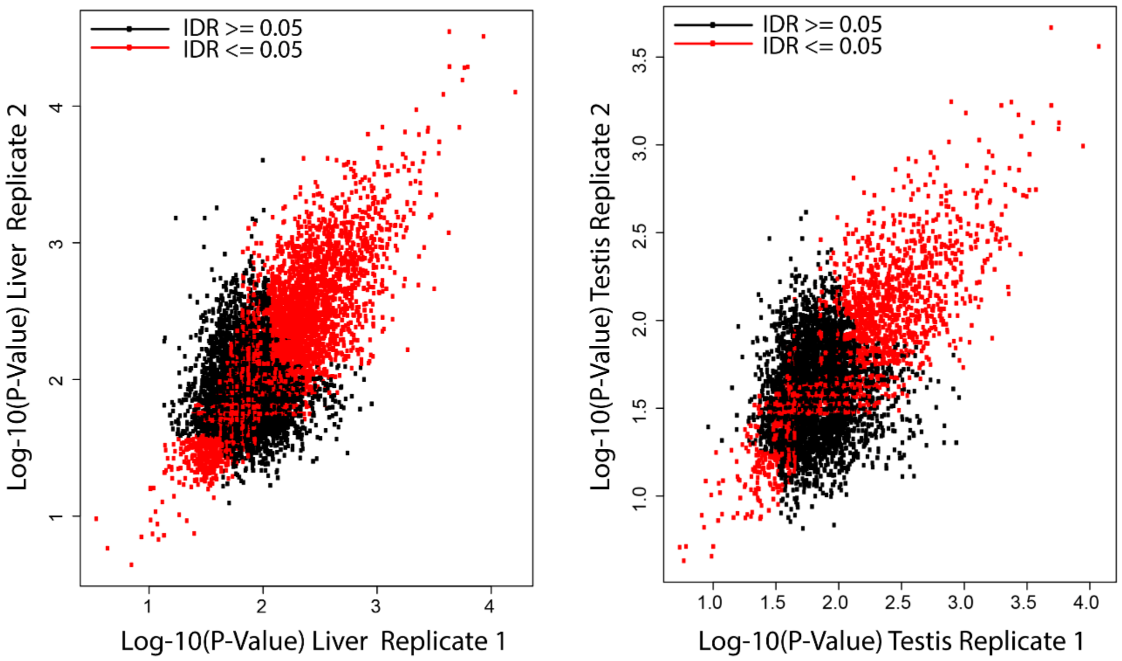


**Supplemental Figure 7. High concordance of chromatin accessibility between peaks identified in two replicates.**

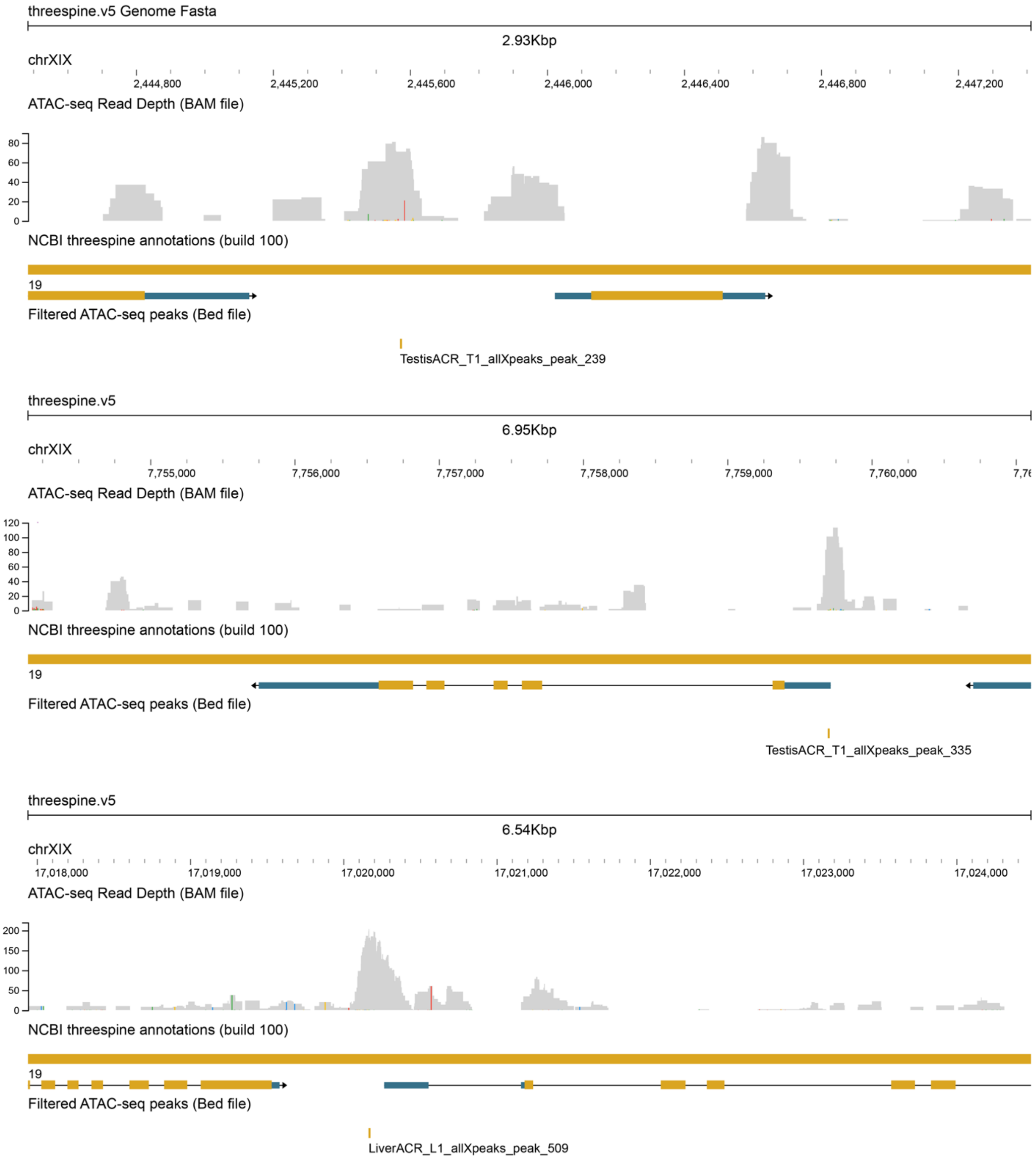


**Supplemental Figure 8. Verification of ATAC-seq proximity to annotated genes.**
